# Cis-Regulatory Element and Transcription Factor Circuitry Required for Cell-Type Specific Expression of FOXP3

**DOI:** 10.1101/2024.08.30.610436

**Authors:** Jennifer M. Umhoefer, Maya M. Arce, Sean Whalen, Rama Dajani, Laine Goudy, Sivakanthan Kasinathan, Julia A. Belk, Wenxi Zhang, Royce Zhou, Sanjana Subramanya, Rosmely Hernandez, Carinna Tran, Nikhita Kirthivasan, Jacob W. Freimer, Cody T. Mowery, Vinh Nguyen, Mineto Ota, Benjamin G. Gowen, Dimitre R. Simeonov, Gemma L. Curie, Zhongmei Li, Jacob E. Corn, Howard Y. Chang, Luke A. Gilbert, Ansuman T. Satpathy, Katherine S. Pollard, Alexander Marson

## Abstract

FOXP3 is a lineage-defining transcription factor (TF) for immune-suppressive regulatory T cells (Tregs). While mice exclusively express FOXP3 in Tregs, humans also transiently express FOXP3 in stimulated conventional CD4+ T cells (Tconvs). Mechanisms governing these distinct expression patterns remain unknown. Here, we performed CRISPR screens tiling the *FOXP3* locus and targeting TFs in human Tregs and Tconvs to discover cis-regulatory elements (CREs) and trans-regulators of FOXP3. Tconv FOXP3 expression depended on a subset of Treg CREs and Tconv-selective positive (TcNS+) and negative (TcNS-) CREs. The CREs are occupied and regulated by TFs we identified as critical regulators of FOXP3. Finally, mutagenesis of murine TcNS- revealed that it is critical for restriction of FOXP3 expression to Tregs. We discover CRE and TF circuitry controlling FOXP3 expression and reveal evolution of mechanisms regulating a gene indispensable to immune homeostasis.

**Highlights:**

- Comprehensive CRISPR maps of CREs and TFs controlling FOXP3 in human Tregs and Tconvs
- Key TFs that control FOXP3 directly occupy and regulate CREs forming TF-CRE circuits
- A previously unknown negative CRE stringently restricts FOXP3 to Tregs in mice

## INTRODUCTION

Lineage-defining factors must be tightly regulated by cis-regulatory elements (CREs) to ensure proper cell type specific expression^1–3^. Indeed, genes encoding these factors are often marked with extended regions of chromatin modifications associated with CRE activity^3,4^. These “super-enhancers” can harbor multiple distinct CREs, each occupied by distinct sets of transcription factors (TFs) and other trans-regulatory factors^3,5–7^. Multiple CRE-TF circuits regulating a key factor allows for robust control of expression that can also respond to multiple signaling inputs^8–10^. While chromatin mapping provides snapshots of candidate CREs and TFs occupying a locus, comprehensive perturbation studies of non-coding sequences and TFs are required to determine which factors are critical to regulate a target gene positively or negatively. Such studies are needed in human immune cells, where closely related cell types can have distinct and even opposing functions in immune homeostasis. Lineage-defining factors enable specialization and shape the balance of inflammatory and immune suppressive cell populations.

CD4+ T cells represent a critical immune compartment comprising closely related, but specialized, subsets of pro-inflammatory cells required to protect against pathogens and suppressive cells required to prevent autoimmunity. Regulatory T cells (Tregs) are a suppressive population of CD4+ T cells that maintain self-tolerance. The TF FOXP3 is a lineage-defining factor in Tregs, and its continued expression is crucial for proper Treg differentiation, suppressive function, and maintenance of cellular identity. In mice, expression of FOXP3 is exclusive to Tregs and serves as a useful marker for Treg identification^11,12^. However, Treg-specific expression of FOXP3 is not conserved in human cells. While human Tregs constitutively express FOXP3 like murine Tregs, human CD4+CD25- conventional T cells (Tconvs) also transiently express FOXP3 upon cellular activation^13–15^. In vitro stimulation via CD3 is sufficient to induce FOXP3 expression in a subset of human Tconvs, and combination with CD28 stimulation and/or exogenous IL-2 further enhances the proportion of FOXP3+ Tconvs in vitro^13,14^. Strong in vitro activation signals are capable of inducing FOXP3 expression in almost all human Tconvs^13,14^. Unlike its function in Tregs, transient expression of FOXP3 in Tconv does not prevent proliferation or expression of pro-inflammatory cytokines, including IL-2 and INF- γ^14,15^. FOXP3 expression in Tconvs has been shown to decrease sensitivity to restimulation-induced cell death^16^, indicating a potential role in modulating activation responses.

Several non-coding elements governing FOXP3 expression in murine Tregs have been characterized in germline deletion mouse models. Four conserved noncoding sequences with FOXP3 enhancer activity in Tregs, named CNS0-3, have been identified. These cis-regulatory elements (CREs) regulate FOXP3 expression in distinct conditions and cell states. CNS0, which lies upstream of the *FOXP3* promoter, regulates IL-2 induced FOXP3 expression during thymic Treg development^17,18^. CNS1 and CNS2 lie within the first intron of *FOXP3*. CNS1 deficiency in mice impairs peripheral induction of FOXP3 and differentiation of induced Tregs (iTregs) from naïve CD4+ T cells^9^. CNS2 (also known as the Treg Specific Demethylated Region, or TSDR) controls heritable maintenance of FOXP3, and its stable activity is dependent on demethylation of CpG dinucleotides within the CRE^9,19^. CNS2 is demethylated in Tregs but near-completely methylated in Tconvs, including naïve CD4+ T cells that are capable of differentiation into iTregs^9,19,20^. Finally, CNS3 lies within the second intron of *FOXP3* and regulates de novo induction of FOXP3 expression in thymic and peripheral Treg development^9^. However, comprehensive characterization of both CREs and TFs in human Tregs remains incomplete.

Mechanisms regulating transient FOXP3 expression in Tconvs, which require functional studies in human cells and not murine models, remain unexamined.

To discover CREs and trans-regulators of FOXP3 expression in human CD4+ T cells, we performed CRISPR screens tiling the *FOXP3* locus and targeting TFs in Tregs and Tconvs. We used a massive pooled CRISPR interference (CRISPRi)-based tiling approach coupled with fluorescence-activated cell sorting (FACS) to inactivate CREs throughout the locus and discover critical elements^7^. Here, CRISPRi tiling screens identified strong Treg and Tconv enhancers of FOXP3 expression at CNS0 in addition to a novel Tconv-specific noncoding sequence (TcNS+) located upstream of CNS0. Validations of these sites additionally identified and fine-mapped a CRE overlapping the long noncoding RNA (lncRNA) *FLICR* that selectively dampens FOXP3 levels in Tregs. Most surprisingly, we discovered a previously unappreciated silencing CRE that dampens FOXP3 in Tconvs (negative Tconv-specific noncoding sequence; TcNS-), especially under resting conditions. CRISPR screens identified TFs required for proper FOXP3 regulation in Tregs and Tconvs, identifying cell type-specific and shared trans-factors that physically interact with FOXP3 CREs including the novel TcNS-. When we did not discover Tconv enhancers that appear active in human but not in mouse Tconvs (which do not express FOXP3), we hypothesized that murine TcNS- may be responsible for total restriction of FOXP3 to Tregs. Indeed, CRISPR disruption of TcNS- caused mouse Tconvs to express FOXP3. Collectively, by systematically mapping CREs and TFs that control a gene critical for immune homeostasis in distinct primary cell subsets, we discover cell-type selective CRE-TF circuits that tightly control expression dynamics. Remarkably, a previously unknown silencer element in the *FOXP3* locus plays a critical role in dampening FOXP3 expression in human Tconvs and strictly blocking expression in murine Tconvs, providing new insight into the evolution of gene control.

## RESULTS

### CRISPRi tiling screen identifies CREs controlling FOXP3 expression

Our initial goal was to create the first comprehensive functional map of FOXP3 CREs in primary human Tregs and Tconvs. We designed a CRISPRi tiling screen with pooled guide RNAs (gRNAs) targeting sites across the *FOXP3* locus (Figure 1A). The gRNA library consisted of ∼15K gRNA spanning a ∼123 kb region from ∼39 kb downstream of the *FOXP3* transcriptional start site (TSS) to ∼85 kb upstream of the TSS. We isolated CD4+CD25^high^CD127^low^ human Tregs and CD4+CD25^low^ human Tconvs from the blood of two healthy donors (Figure S1A), stimulated cells, and delivered dCas9-ZIM3 CRISPRi machinery and the gRNA library via lentivirus. Cells were restimulated for 48 hours to induce FOXP3 expression (Figure S1B), and cells were FACS-sorted into bins of high (top ∼25%) and low (bottom ∼25%) FOXP3 expression (Figure S1C). We achieved sufficient coverage to identify CREs involved in both maintenance and repression of FOXP3 expression (Figure 1B, Figure S1D-E), and results were largely consistent between donors (Figure S1F).

**Figure 1.**
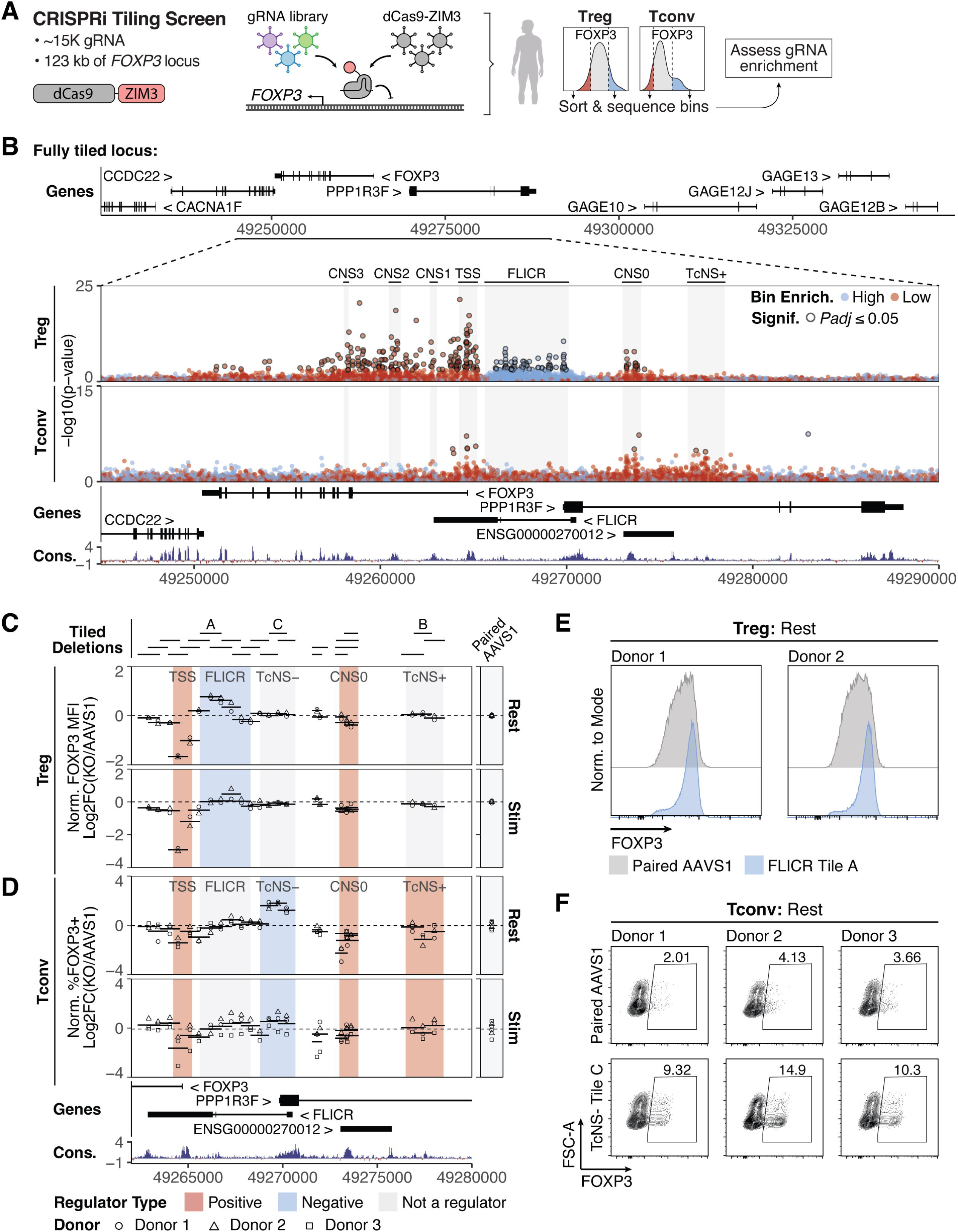
CRISPRi tiling screen identifies FOXP3 CREs in Tregs and Tconvs. (A) Schematic depicting CRISPRi-based screens for FOXP3 CREs. (B) Top, full assayed region in CRISPRi *FOXP3* locus-tiling screen. Bottom subplot, -log10(p- value) of gRNA enrichment in FOXP3 high vs. low FACS bins in Treg and Tconv. Blue, gRNA enriched in FOXP3 High FACS bin. Red, gRNA enriched in FOXP3 Low FACS bin. Outlined, adjusted p-value ≤ 0.05. Treg CNS0 and CNS2, *FOXP3* TSS, the FLICR region, and TcNS+ are highlighted in grey. Treg, *n* = 2 donors; Tconv, *n* = 2 donors. Conservation track depicts vertebrate PhyloP 100-way conservation. (C and D) Arrayed validation of *FOXP3* TSS, FLICR, CNS0, and TcNS+ region-associated CRISPRi-responsive elements with paired Cas9 RNPs plotted along the *FOXP3* locus at 0 hours (Rest) and 48 hours (Stim) post-stimulation in Tregs (C) and Tconv (D). Locations of tiled deletions are indicated by line segments in (C) and (D). Lines indicate log2 fold change in mean FOXP3 MFI in Tregs (C) or %FOXP3+ Tconvs (D) in deletions vs. donor-specific paired *AAVS1* gRNA controls. Points indicate individual replicates of tiled deletions (Treg *n* = 2 donors; Tconv *n*= 3 donors). Positive CREs for each cell type are highlighted in red, negative CREs in blue, and non-regulatory regions in gray. (E) Flow plots of FOXP3 expression upon deletion of Tile A and an *AAVS1* control in resting Tregs. Tile letter corresponds to Tiled Deletions in (C, D). (F) Representative flow plots depicting FOXP3 expression with Tile C and paired *AAVS1* deletions in resting Tconv. Numbers indicate %FOXP3+ cells. Tile letter corresponds to Tiled Deletions in (C, D).

In Tregs, the TSS of *FOXP3* was the most responsive to CRISPRi tiling, resulting in significant gRNA enrichment in the FOXP3 low bin. Numerous gRNAs across the first 8 kb of the *FOXP3* gene body were enriched in the FOXP3 low bin, including gRNAs mapping to CNS1, 2, and 3. CRISPRi screening also revealed that activity at CNS0 is required to maintain FOXP3 expression in human Tregs, which was striking because its individual role in FOXP3 maintenance in mature murine Tregs has been ambiguous^17,18^. In addition to mapping conserved enhancer elements in the locus, we also identified a Treg-specific 4.6 kb FOXP3-repressive element 1.1 kb upstream of the *FOXP3* TSS that maps to *FLICR*, a lncRNA transcript described in mice^21^. Previous work has shown that *Flicr* is specifically expressed in Tregs and acts in cis to mildly repress *Foxp3* expression^21^ (Figure S1G); we now demonstrate *FLICR* is also functional in human Tregs.

The landscape of functional elements in the locus identified by CRISPRi in human Tconvs was distinct from what we observed in Tregs. CNS1, 2, and 3 did not appear to be essential for FOXP3 induction in Tconvs (Figure 1B). The promoter and CNS0 were critical for FOXP3 expression in human Tconvs in addition to Tregs, and an additional 2 kb partially-conserved non-coding sequence (termed here positive Tconv Non-Coding Sequence, TcNS+) approximately 11.8 kb upstream of the *FOXP3* TSS emerged as a selective regulator of FOXP3 induction in Tconvs (Figure 1B). From these screens, we conclude that FOXP3 expression in human Tregs is maintained by CNS0 and enhancers within the gene body and suppressed by an upstream CRE mapping to the lncRNA *FLICR*. FOXP3 induction in Tconvs depends on a set of CREs distinct from the well-characterized Treg enhancers including TcNS+, which displays selective function in Tconvs.

To validate the results of the CRISPRi screens and quantify effects of CRE perturbation on FOXP3 expression, we designed an orthogonal CRISPR nuclease (CRISPRn) deletion strategy using paired Cas9 RNPs to excise individual candidate CREs of interest. Using this strategy, we generated 28 different ∼1 kb deletions across the *FOXP3* TSS, *FLICR*, CNS0, and TcNS+ in resting and stimulated human Tregs and Tconvs (Figure S2A-B). For each deletion, we measured FOXP3 median fluorescence intensity (MFI; Tregs) and the percent of FOXP3+ cells (Tconvs) for each targeted deletion and for *AAVS1*-targeted control cells (Figure 1C-D).

Deletions in Tregs confirmed that the TSS and CNS0 were required to maintain FOXP3 levels in both resting and stimulated Treg (Figure 1C). They also confirmed that *FLICR* tunes down FOXP3 levels in human Tregs. CRISPR deletions fine-mapped functional sequences overlapping a shorter *FLICR* isoform that is the predominant transcript detected in long-read sequencing from human PBMCs (Figure S2A). In particular, deletion of a region overlapping with the TSS of the shortened isoform resulted in dramatic up-regulation of FOXP3 in resting Tregs (Tile A, Figure 1C, E, Figure S2A). These results define critical functional CREs upstream of the *FOXP3* TSS in human Tregs, where CNS0 contributes to FOXP3 maintenance and the *FLICR* region acts to limit FOXP3 expression.

In Tconvs, deletions targeting the TSS or CNS0 decreased the percentage of FOXP3+ cells (Figure 1D). Deletion of one region of TcNS+ led to a decreased percentage of FOXP3+ cells in resting Tconvs and a mild decrease in activated Tconvs (Tile B, Figure 1D). Interestingly, in resting Tconvs, deletion of genomic elements overlapping with the promoter and TSS of the neighboring gene *PPP1R3F* resulted in a large increase in the percent of FOXP3+ Tconvs (Tile C, Figure 1D, F). This effect was more modest in stimulated Tconvs, the condition used in the pooled CRISPRi screen. KO gRNAs targeting the coding region of *PPP1R3F* did not increase FOXP3 expression in resting or stimulated Tconvs, indicating non-coding sequences near the *PPP1R3F* promoter are likely responsible for FOXP3 regulation (not the *PPP1R3F* gene product itself; Figure S2C). Furthermore, *FLICR* is not expressed in Tconvs, so these non-coding deletions are unlikely to act via the lncRNA (Figure S1G). We termed this non-coding element in the *PPP1R3F* promoter the negative Tconv Non-Coding Sequence (TcNS-), due to its function in repressing FOXP3 expression in Tconvs. As a whole, these results confirm that CNS0 and TcNS+ are both required for normal FOXP3 expression in human Tconvs, and, surprisingly, TcNS- limits FOXP3 expression in Tconvs, particularly notably in resting Tconvs.

### Chromatin hallmarks of enhancer activity at CNS0, TcNS+, and TcNS-

We next measured the chromatin state of the CREs in resting and stimulated Tregs and Tconvs. We analyzed ATAC-seq in unstimulated and stimulated Tregs and Tconvs^22^ to parse the chromatin accessibility landscape at the FOXP3 locus. In unstimulated cells, a highly accessible peak overlapped TcNS- in both cell types, and a smaller peak mapped to CNS0, especially in Tregs (Figure 2A, Figure S3A). However, upon stimulation, accessibility increased at TcNS+ and CNS0 in both cell types (Figure 2A, Figure S3A). Stimulation-responsive accessibility at TcNS+ and CNS0 was particularly notable in Tconvs, mirroring the stimulation-responsive induction of FOXP3 in human Tconvs.

**Figure 2.**
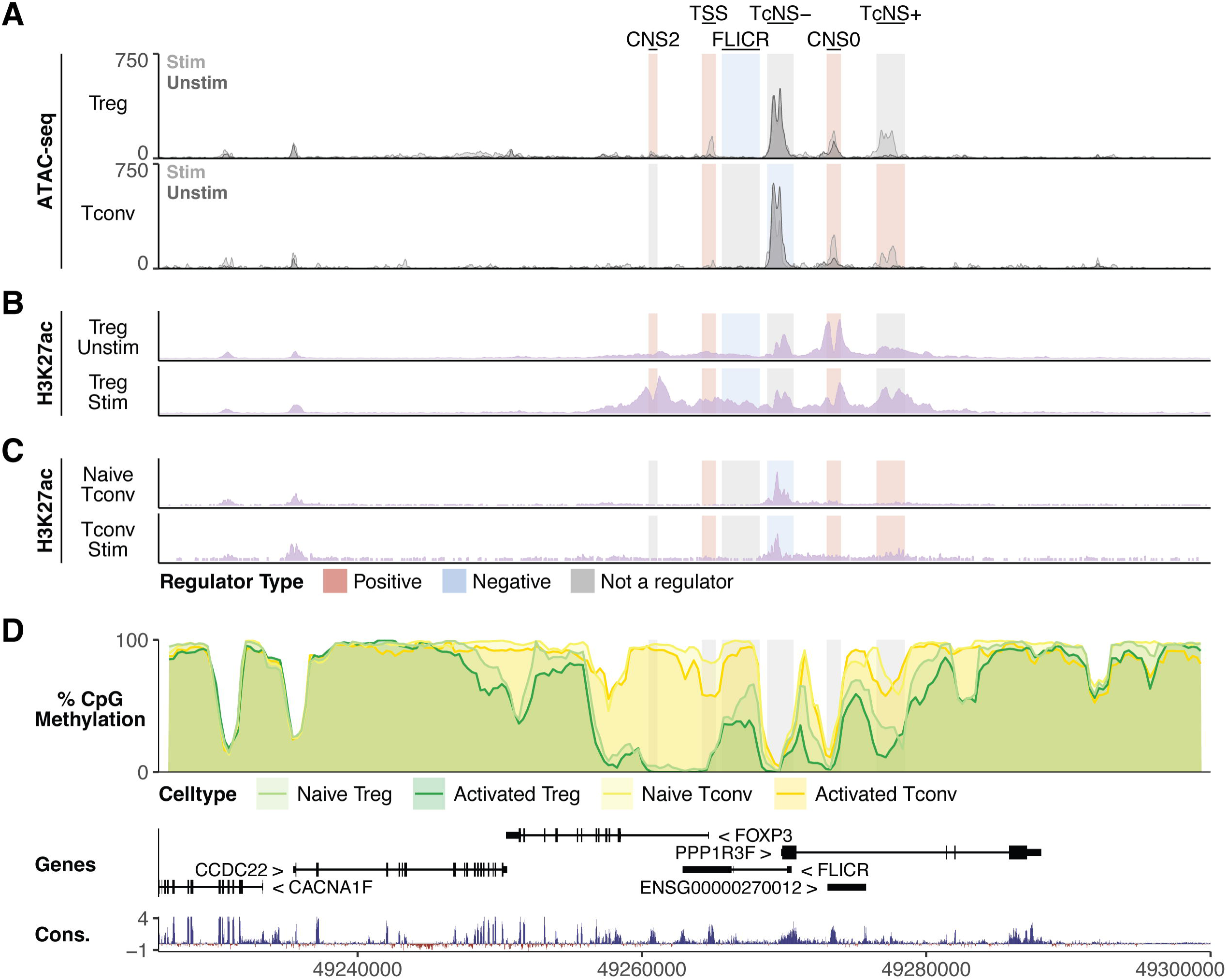
Positive and negative FOXP3 CREs map to active chromatin. (A) ATAC-seq in unstimulated (dark gray) and stimulated (light gray) Treg and Tconv^22^. Positive CREs for the indicated cell type are highlighted in red, negative CRE in blue, and non- regulatory regions in gray. (B and C) ChIP-seq for H3K27ac in unstimulated (top) and stimulated (bottom) Tregs (B) and Tconvs (C). ChIP-seq is auto-scaled to the locus shown. (D) Post-bisulfite-conversion adapter tagging sequencing (PBAT-seq) percent methylation of CpGs at the *FOXP3* locus in naïve and activated Tregs (green) and naïve and activated Tconvs (gold) from a representative donor. PBAT-seq data is plotted as the mean percent methylation of 1500 bp bins with 300 bp step sizes across the locus shown.

To assess chromatin marks associated with enhancer activity further, we conducted ChIP-seq in Tregs and analyzed ChIP-seq in Tconv for H3K27ac^1^, a marker of active enhancers. Tregs showed strong H3K27ac signal across the *FOXP3* gene body and upstream region, with notable peaks at CNS2, CNS0, and TcNS+ (Figure 2B), consistent with enhancer activity at CNS2 and CNS0. Interestingly, H3K27ac was observed at TcNS- in both Tregs and Tconvs despite functionally serving to dampen FOXP3 expression in Tconvs (Figure 2B-C). This observation accords with prior reports that CREs that function as silencers can also be marked with H3K27ac^23^.

Given the well-known role of DNA methylation in regulating FOXP3 expression, we analyzed post-bisulfite-conversion adapter tagging sequencing (PBAT-seq) in naïve and activated Tregs and Tconvs^24^ to assess CpG methylation at CREs. Strikingly, CpGs in Tregs and Tconvs were highly hypomethylated at TcNS- and CNS0. CpGs within TcNS+ were partially hypomethylated in Tregs and also to a lesser extent in Tconvs (Figure 2D). Taken together, these data indicate CNS0 and TcNS+ are stimulation responsive elements with histone modifications and DNA methylation patterns associated with enhancer activity. TcNS-, which serves as a silencer of FOXP3 in Tconvs, is unmethylated in CD4+ T cells with marks of active chromatin.

### CRISPR KOs define critical TFs for FOXP3 expression

To identify TFs required for FOXP3 expression that may bind to FOXP3 CREs, we performed pooled CRISPRn knockout screens in primary human Tregs and Tconvs using a library of 6000 gRNAs targeting 1349 human TFs, chromatin modifiers, and immune genes (Figure 3A; Figure S4A)^25,26^. 48 hours after restimulation, cells were sorted via FACS into bins of high (top ∼25%) and low (bottom ∼25%) FOXP3 expression and sequenced to assess gRNA enrichment (Figure 3A, Figure S4B). We identified 22 positive and 9 negative regulators of FOXP3 in human Tregs, and 23 positive and 15 negative regulators in human Tconvs (FDR ≤ 0.05; Figure 3B-C). We achieved sufficient coverage across donors and cell types (Figure S4C-D), and high reproducibility of significant hits was observed between donors (Figure S4E-F). Additionally, individual gRNAs targeting key gene regulators were enriched consistently in FOXP3 high or low FACS bins in both cell types (Figure 3D).

**Figure 3.**
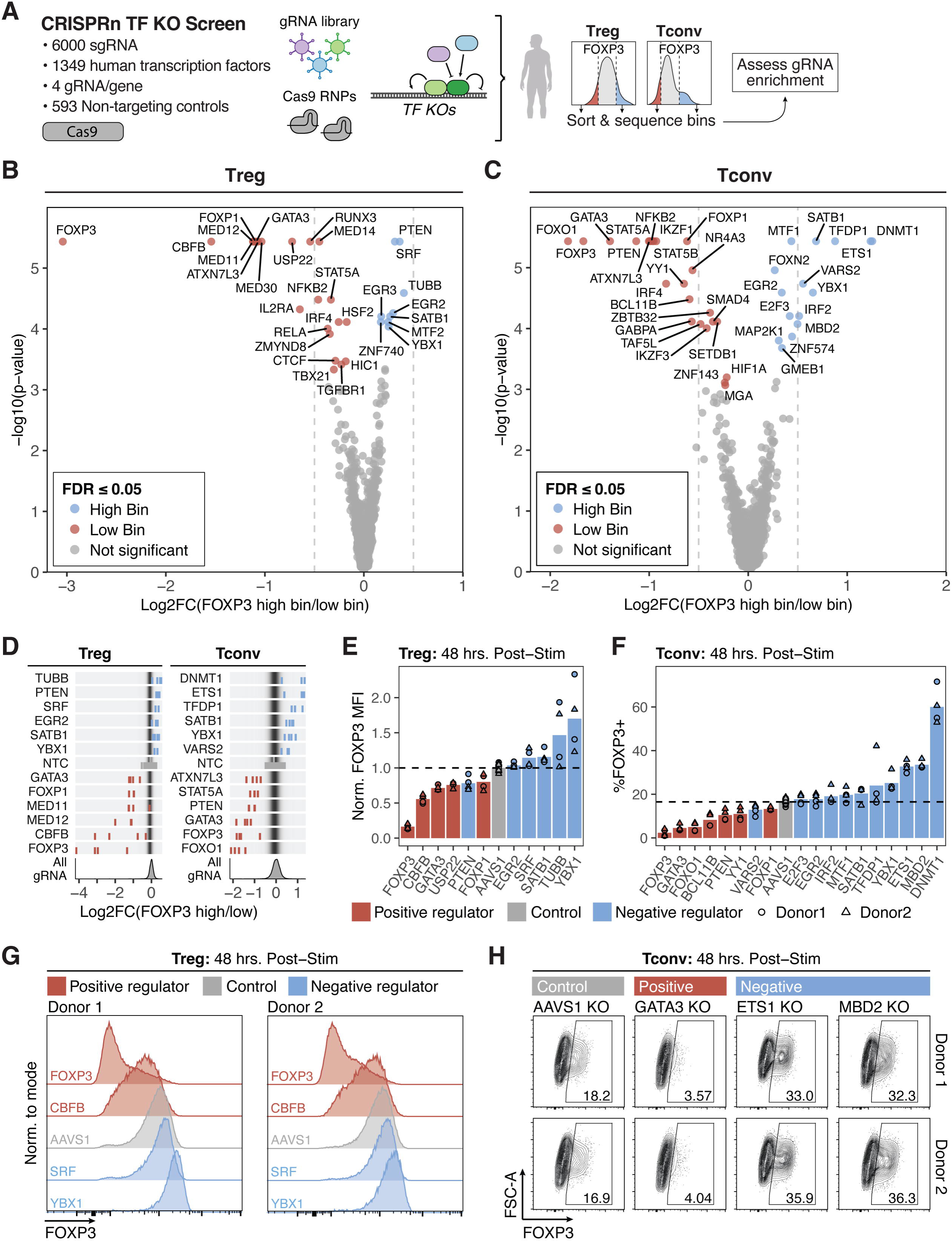
CRISPRn TF screens identify trans-regulators of FOXP3 in Tconv. (A) Schematic depicting CRISPRn-based screens for FOXP3 trans-regulators in Tregs and Tconvs. (B and C) Volcano plots of log2 fold change of gene gRNA enrichment in FOXP3 high vs. low FACS bins versus -log10 of p-value in Tregs (B) and Tconvs (C). Color of points indicates significance (FDR ≤ 0.05). Blue, significantly enriched in FOXP3 high FACS bin. Red, significantly enriched in FOXP3 low FACS bin. Gray, not significant (Treg, *n* = 2 donors; Tconv *n* = 3 donors). (D) Top, log2 fold change of individual gRNA enrichment in FOXP3 high vs. low FACS bins for the top 12 significant FOXP3 positive and negative regulators in Tregs (left) and Tconvs (right, FDR ≤ 0.05; *n* = 2 Treg donors, *n* = 3 Tconv donors). gRNAs corresponding to the labelled significant gene are colored blue (gene enriched in FOXP3 high bin), red (gene enriched in FOXP3 low bin), or gray (NTC, non-targeting control gRNA). Bottom, distribution of gRNA in screen. (E and F) Arrayed validation of top FOXP3 positive and negative regulators at 48 hours post-stimulation in Tregs (E) and Tconvs (F). Color indicates the directional effect in the screen (*n* = 2 donors x 2 gRNAs per target or 6 gRNAs targeting *AAVS1* control). (G) Selected flow plots of FOXP3, CBFB, *AAVS1*, SRF, and YBX1 KO in stimulated Tregs. (H) Selected flow plots of *AAVS1*, GATA3, ETS1, and MBD2 KO in Tconvs at 48 hours post-stimulation. Numbers indicate %FOXP3+ cells.

In Tregs, several previously described positive regulators with importance in murine Tregs emerged among hits in the screen, including CBFB, GATA3, ATXN7L3, USP22, and STAT5A^17,27–30^ (Figure 3B, Figure S4G). Notably, Treg FOXP3 expression depended on multiple components of the mediator complex (MED11, MED12, MED14, and MED30) and was dampened by SATB1 and SRF (Figure 3B, Figure S4G). In addition, we identified YBX1, which has reported RNA and DNA binding activity, as a previously unappreciated negative regulator of FOXP3 in human Tregs absent from murine studies (Figure 3B, Figure S4G). In Tconvs, a subset of positive regulators were shared with Tregs, including GATA3^28^, STAT5A/STAT5B^17,30^, and ATXN7L3^29^ (Figure 3C), which until now have not been shown to influence FOXP3 expression in human Tconvs. The human Tconv screen also revealed strong negative regulators of FOXP3, including methylation-associated factors (e.g., DNMT1 and methyl-CpG binding domain protein, MBD2), regulators shared with Tregs (e.g., YBX1, SATB1 and EGR2), and regulators that were selectively found in our Tconv screen (e.g., TFDP1 and ETS1) (Figure 3C).

Interestingly, loss of the maintenance DNA methyltransferase DNMT1 also causes FOXP3 induction in murine Tconvs and CD8 T cells (especially in the presence of T cell receptor stimulation), underscoring a critical role for DNA methylation in FOXP3 expression^31,32^.

Together, these studies revealed the distinct sets of TFs that regulate the same gene (FOXP3) in different T cell populations.

We next validated the effects of individual TF KOs. For KOs of top candidate TFs that promoted or repressed FOXP3 in the pooled screen, we measured FOXP3 MFI (Tregs) or the percentage of FOXP3+ cells (Tconvs, Figure S5A) in resting and stimulated (48 hours post-stimulation) states via flow cytometry. We achieved high editing efficiency of KOs as assessed by targeted amplicon sequencing (mean percentage modified reads of 82.3% in Tregs, 90.5% in Tconvs; Figure S5B).

KO of regulators in Tregs and Tconvs largely replicated the directionality of FOXP3 changes observed in the screen (Figure 3E-H, Figure S5C-D). In Tregs, YBX1 was the strongest negative FOXP3 regulator, with KO significantly increasing FOXP3 MFI in both resting and stimulated Tregs over *AAVS1*-targeting controls (Figure 3E, G, Figure S5C). In Tconvs, GATA3 KO resulted in a large reduction in the percentage of FOXP3+ cells at 48 hours post-stimulation, decreasing the percentage of FOXP3+ cells nearly to levels of FOXP3 KO (Figure 3F, H).

Among negative regulators, KO of TFDP1, ETS1, DNMT1, YBX1, and MBD2 resulted in the largest increases in FOXP3 expression in Tconvs (Figure 3F, H, Figure S5D). High baseline levels of FOXP3 expression in Tregs and low baseline expression in Tconvs enabled enhanced detection of positive regulators in the Treg screen and negative regulators in the Tconv screen. Arrayed KO of Treg or Tconv regulators in the opposite cell type revealed additional FOXP3- suppressive activity of TFDP1 and DNMT1 in Tregs and FOXP3-promoting activity of CBFB and USP22 in Tconvs (Figure S5E-F). Collectively, CRISPRn screening and RNP-based validation identified known and novel regulators of FOXP3 expression in Tregs, including unappreciated negative regulators of FOXP3, and comprehensively characterized FOXP3 regulators in Tconvs, which were previously unknown.

### Trans-regulators bind to TcNS-, CNS0, and TcNS+ to control FOXP3 expression

We tested if TFs identified as critical regulators of human FOXP3 operate directly on CREs within the *FOXP3* locus. We conducted ChIP-seq for select regulators in Tregs and analyzed available regulator ChIP-seq datasets in Tregs, T cells, and T cell lines^33,34,43,35–42^. In Tregs, FOXP3, STAT5, and HIC1 (positive regulators of FOXP3) primarily bound positive CRE regions, including FOXP3 binding at CNS0, STAT5 binding at CNS0 and CNS2, and HIC1 binding at CNS0 and the TSS (Figure 4A). Interestingly, STAT5 and HIC1 additionally bound TcNS+, even though TcNS+ is seemingly dispensable for FOXP3 maintenance in mature Tregs. Negative regulators additionally bound positive CREs, including SRF binding at CNS0 and SATB1 binding at CNS0, the TSS, and weak binding at CNS2 (Figure 4A), suggesting a balancing effect of positive and negative regulators at these enhancers. YBX1 ChIP-seq yielded no detectable peaks at the FOXP3 locus (Figure S6A), suggesting that YBX1 may operate as a strong negative regulator of FOXP3 indirectly or at the post-transcriptional level through its RNA-binding activity. Notably, Tconv positive regulators, GATA3, STAT5B, and BCL11B bound CNS0, and STAT5B also bound TcNS+ (Figure 4B). Motif analysis using predicted TF binding motifs suggested additional potential direct interactions of regulator TFs at FOXP3 CREs in Tregs and Tconvs (Figure S7A-B). In short, ChIP-seq analysis suggests that multiple factors promoting FOXP3 expression can bind directly to CNS0 in Tregs and CNS0 and TcNS+ in Tconvs.

**Figure 4.**
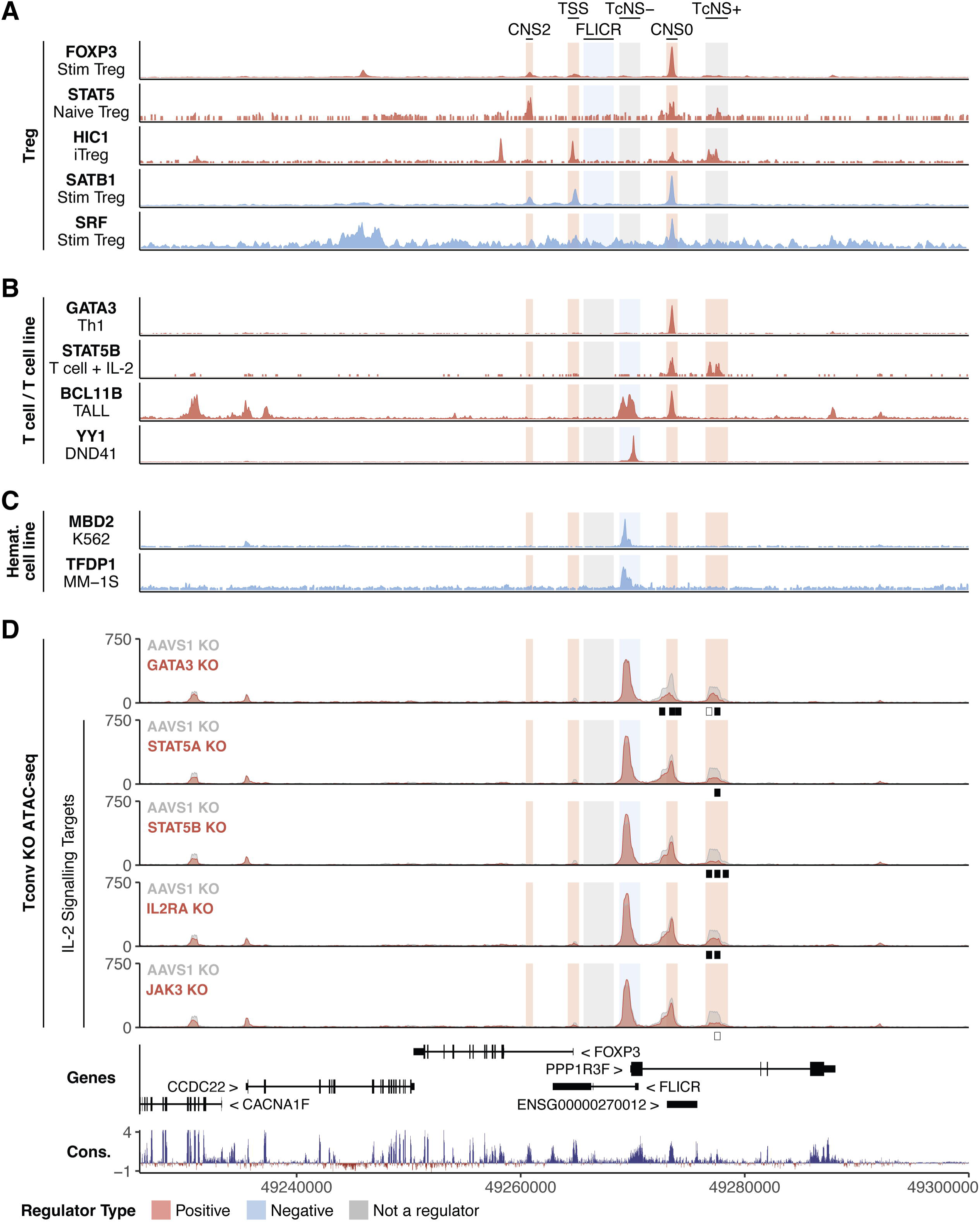
KO of positive and negative trans-regulators change accessibility at CNS0 and TcNS+. (A) ChIP-seq in Tregs targeting Treg positive (red) and negative (blue) trans-regulators of FOXP3. CREs are colored based on the effect of the element. Red, positive regulator; blue, negative regulator; gray, not required for regulation. ChIP-seq is auto-scaled to the locus shown. (B and C) Publicly available ChIP-seq datasets in indicated T cells or T cell-based cell lines (B) or hematopoietic cell lines (C) targeting Tconv positive (red) and negative (blue) trans-regulators of FOXP3. CREs are labelled and scaling conducted as described in (A). (D) Effect of trans-regulator KO on DNA accessibility^26^ at the *FOXP3* locus. Bars beneath plots indicate significantly differentially accessible peaks in KOs vs. *AAVS1* controls (White, *Padj* ≤ 0.1; Black, *Padj* ≤ 0.05). For each KO comparison, *n* = 2-3 donors x 1 KO gRNA and 7-8 *AAVS1*-targeting gRNAs. Red, accessibility with indicated positive regulator KO; gray, accessibility in *AAVS1* controls. In CRISPRn TF KO screens, IL2RA and JAK3 were just above an FDR cutoff of 0.05.

We also searched for factors that directly occupy TcNS-, the CRE with silencer activity in Tconvs. Strikingly, binding motifs for numerous negative FOXP3 trans-regulators nominated by our TF screens in Tconvs clustered within TcNS-, including EGR2, ETS1, DNMT1, TFDP1, and MBD2 (Figure S7B). Two of these negative regulators, MBD2 and TFDP1, can bind strongly and selectively to TcNS- in hematopoietic cell lines (Figure 4C). Beyond these enhancers, multiple regulators bound to a site near the *CCDC22* TSS (Figure 4B, Figure S6B). We also noted that a subset of positive regulators can bind to TcNS- including BCL11B and YY1, which is associated with maintenance of enhancer-promoter loops and activating and repressive chromatin modifiers^44,45^ (Figure 4B), again suggesting a balance of positive and negative factors at the element. Interestingly, binding motifs for ZBTB32, a positive regulator with reported chromatin-repressive activity^46,47^, also aligned proximal to TcNS- (Figure S7B). Taken together, several TF regulators of FOXP3 bind directly to TcNS-, including TFs that serve to dampen FOXP3 levels in Tconvs.

We next assessed if these TFs are also required to regulate the chromatin state of CREs in human Tconvs. We reasoned that TF KOs paired with ATAC-seq may reveal altered accessibility at the FOXP3 locus. Therefore, we analyzed published datasets from our group that paired TF KOs with ATAC-seq in CD4+CD25- Tconvs^26^. Among the 11 FOXP3 regulator TFs included in the KO ATAC-seq dataset, GATA3, STAT5A, and STAT5B KO resulted in highly differentially accessible peaks in the *FOXP3* locus (Figure 4D). KO of positive regulator GATA3 caused decreased accessibility at TcNS+ and CNS0, which aligned with a distinct GATA3 ChIP-seq peak at CNS0 (Figure 4B, D). Therefore, GATA3 likely maintains FOXP3 expression in part via direct maintenance of CNS0 accessibility.

STAT5A KO and STAT5B KO caused decreased accessibility at TcNS+ (Figure 4D), which also aligned with STAT5B peaks at this site in IL-2 treated T cells (Figure 4B). Previous studies have shown that IL-2, in combination with anti-CD3 stimulation, boosts Tconv expression of FOXP3 compared to anti-CD3 stimulation alone^14^. Changes in accessibility at TcNS+ with STAT5 KOs nominate TcNS+ as a CRE that may contribute to IL-2-responsive enhancement of FOXP3 expression. We thus assessed differential accessibility resulting from KO of IL2RA or JAK3, critical components of the IL-2 signaling pathway that fell just above our screening significance cutoff of FDR ≤ 0.05 (Figure S8A). IL2RA and JAK3 KO similarly resulted in decreased accessibility at TcNS+ (Figure 4D; note, JAK3 KO *Padj* = 0.066), highlighting TcNS+ as an IL- 2 pathway responsive CRE of FOXP3 in Tconvs. Taken together, these data indicate numerous trans-regulators interact directly with FOXP3 CREs, including GATA3 and STAT5, which maintain accessibility at FOXP3 enhancer elements CNS0 and TcNS+.

### TcNS- restricts FOXP3 expression to Tregs in mice

Mechanisms dictating the marked difference in FOXP3 expression dynamics between human and murine Tconvs are unknown. Systematic elucidation of CREs required for FOXP3 regulation in human Tconvs provides an opportunity to identify CRE sequence evolution that may underlie divergent expression patterns. We initially hypothesized that TcNS+, which is only partially conserved between mouse and human, may permit FOXP3 induction in human Tconvs. However, we found evidence of H3K27ac at both TcNS+ and CNS0 in murine naïve CD4+ T cells, consistent with enhancer activity at these sites despite the lack of FOXP3 expression in these cells (Figure 5A). Furthermore, these murine sites are occupied by positive regulators of FOXP3; GATA3 binds at CNS0, and STAT5 binds at CNS0 and TcNS+, mirroring binding observed in human Tconvs (Figure 5A). These findings suggest that active chromatin at CNS0 and TcNS+ is not sufficient to activate FOXP3 in murine Tconvs.

**Figure 5.**
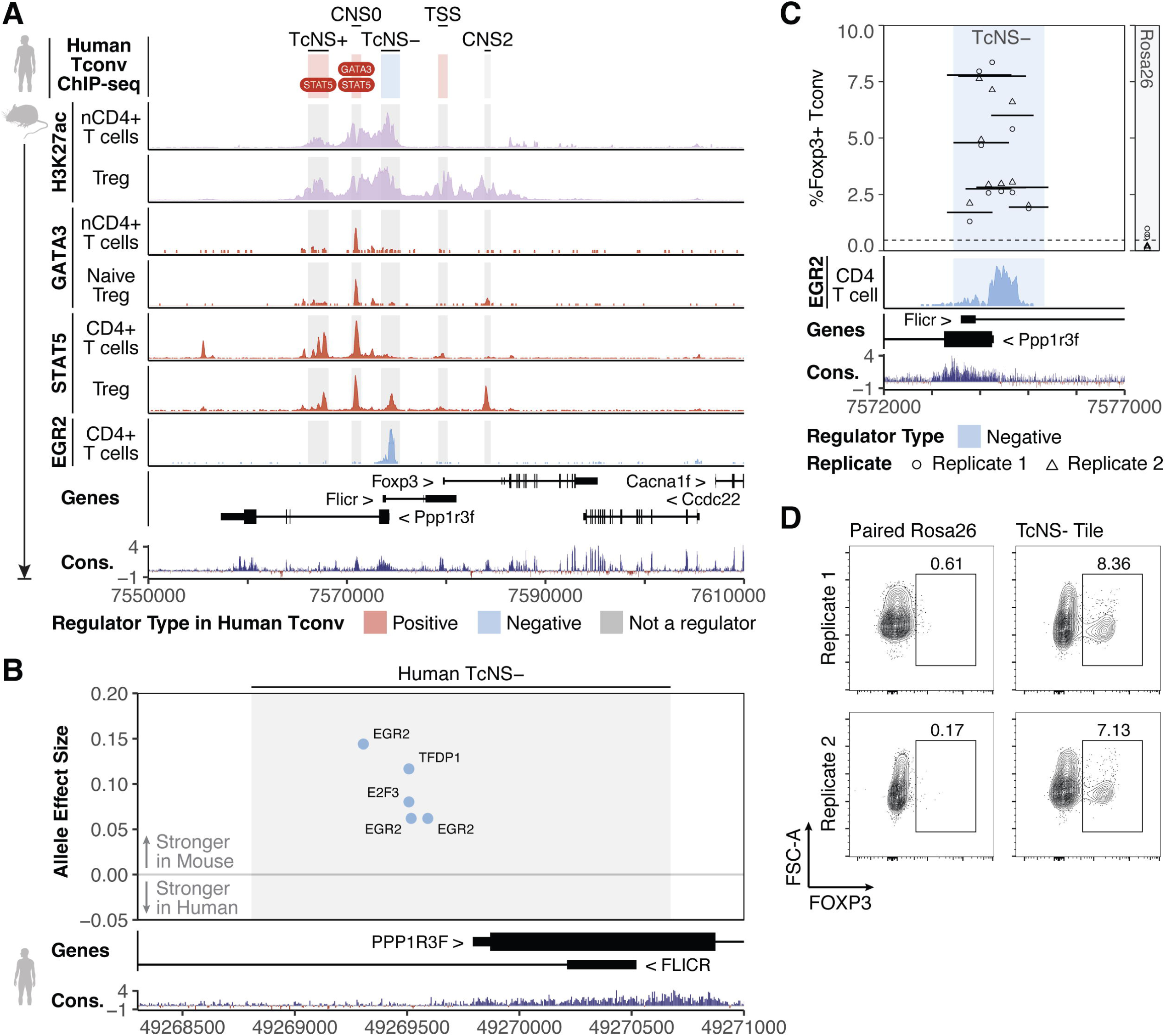
TcNS- suppresses FOXP3 expression in murine Tconv. (A) Top, schematic showing location of GATA3 and STAT5 ChIP-seq in human Tconvs mapped to homologous regions in the murine genome. Below, ChIP-seq for H3K27ac (purple), GATA3, STAT5, and EGR2 in murine Tregs and CD4+ T cells. ChIP-seq is auto-scaled to the locus shown. (B) Allele effect size of TF binding motifs between human (reference allele) and mouse (alternative allele) TcNS-, plotted along the human locus. Allele effect size indicates the difference in motif scoring between the reference and alternate allele (alternate – reference). Only motifs corresponding to human FOXP3 trans-regulators are shown. (C) *Top*, percent FOXP3+ Tconvs with TcNS- tiled deletions. Dashed line indicates the mean %FOXP3+ cells in *Rosa26* controls, shown on the right. Locations of tiled deletions are indicated by line segments, which represent mean %FOXP3+ Tconvs. Points indicate individual replicates of tiled deletions (*n* = 2 replicates). Bottom, EGR2 ChIP-seq in murine CD4+ T cells. ChIP-seq is auto-scaled to the locus shown. (D) Flow plots depicting %FOXP3+ Tconvs with deletion of a segment in *Rosa26* or a tiled deletion at TcNS-.

Similarities in CNS0 and TcNS+ chromatin profiles in human and murine Tconvs led us to a revised hypothesis. Perhaps TcNS- could also have a silencing effect on FOXP3 in murine

Tconvs, which could be stronger than in humans, fully blocking FOXP3 induction in murine Tconvs. We assessed how sequence changes between human and mouse TcNS- are predicted to alter motif affinity of key TFs. By scanning for binding motifs of FOXP3-repressing TFs characterized in human Tconvs, we identified a cluster of EGR2, TFDP1, and E2F3 motifs with stronger predicted affinity in murine TcNS- compared to the homologous human TcNS- sequence (Figure 5B, Figure S9A, Figure S10A-B). Notably, EGR2–a negative FOXP3 regulator in human Tconvs–directly occupies this site in murine Tconvs (Figure 5A). These findings suggested TcNS- could act as a strong FOXP3 silencer in murine Tconvs.

To test if the silencing activity of the murine TcNS- sequence is critical for restricting FOXP3 expression to Tregs, we performed CRISPR deletions of TcNS- in murine Tconv. Briefly, we introduced Cas9 deletions with paired RNPs tiled across TcNS- in murine Tconvs and assessed changes in FOXP3 expression 9 days post-initial stimulation. Paired and individual gRNAs targeting the *Rosa26* safe-harbor locus were used as a control. Indeed, deletion of TcNS- induced FOXP3 expression in murine Tconvs under resting conditions (Figure 5C-D). Several of the most efficacious TcNS- deletions overlapped with a less conserved non-coding region of TcNS- upstream of the *Ppp1r3f* TSS (Figure 5C), where an EGR2 ChIP-seq peak and motif resides. We used individual CRISPRn gRNAs to test requirements of specific sequences in this region for effects on FOXP3 repression (Figure S10A-B). In parallel, we tested the effects of knocking out individual TFs with predicted binding. No individual KO of FOXP3-repressing TFs was sufficient to activate FOXP3 in resting Tconv, perhaps due to redundancy among TF family members (Figure S10C). Yet, mutations introduced by a single gRNA targeting a site in the EGR2 motif were sufficient to induce FOXP3 in resting murine Tconvs, albeit not as strongly as the larger TcNS- deletions (Figure S10C). Collectively, mutational assays in Tconvs indicate that murine TcNS- is required for the restriction of FOXP3 expression to Tregs observed in mice.

The human TcNS- sequence also represses FOXP3 expression in Tconvs, albeit less stringently as transient induction of FOXP3 is possible when human Tconvs are stimulated provided that both TcNS+ and CNS0 are active.

## DISCUSSION

Here, we perform CRISPR screens in primary human CD4+ T cells to decipher the regulatory logic of how distinct cell types control FOXP3 expression via defined TF-CRE circuits. We discovered a critical role for CNS0 in both human Tregs and Tconvs and additionally discovered a Tconv-selective enhancer, TcNS+. A region corresponding to the lncRNA *FLICR* restricted FOXP3 levels in Tregs. Most surprisingly, a novel CRE, TcNS-, was identified as a silencer that serves to dampen FOXP3 levels in Tconvs. Subsets of CRISPR-nominated TFs that regulate FOXP3 levels in Tregs and Tconvs were confirmed to directly bind to CREs, including several key TFs that control accessibility of positive FOXP3 CREs and several others that repress FOXP3 induction in Tconvs and can bind to TcNS-.

FOXP3 is critical for Treg development and function, and, accordingly, it is maintained robustly in Tregs by numerous CREs. Many of these CREs comprise a super-enhancer with broad H3K27ac across the locus in both mouse and human. Prior studies in mice have demonstrated how this highly active region provides robustness; perturbation of individual enhancers, such as CNS0 or CNS3 alone, causes no or mild disruption of FOXP3 expression, while simultaneous targeting of multiple elements markedly destabilizes FOXP3 expression, especially during development^17,18^. Here, CRISPRi allowed us to define CREs individually required to share maintenance of FOXP3 levels in human Tregs. CNS0 activity was required for full maintenance of FOXP3 expression; however, the effects of disrupting this sequence were relatively modest in arrayed experiments and did not completely ablate FOXP3 expression. Targeting multiple CREs simultaneously may reveal stronger effects on FOXP3 levels.

Our functional genomic studies of FOXP3 induction in human Tconvs provided a foothold to investigate the evolutionary switch that allows human Tconvs to express FOXP3 in this context where it is prevented in mice. Despite our initial expectations, we could not find evidence for a human enhancer that is not active in murine cells driving FOXP3 expression in stimulated Tconvs. This led to the surprising discovery that a silencing CRE, TcNS-, has evolved between mice and humans and that the murine version of this element plays a critical role in strictly limiting FOXP3 expression to Tregs in mice. Murine studies over the past several decades have highlighted FOXP3 as a transcription factor exclusively expressed in Tregs and required for the development and function of these cells. We demonstrated that this cell type restriction of FOXP3 to Tregs is lost in the absence of TcNS-. Binding sites for key FOXP3 regulating TFs in this element differ between mouse and humans. Disruption of even one consensus binding site with a single gRNA was sufficient to disrupt the normal expression pattern of murine Tregs.

Much work in recent years has focused on the enhancer code that drives context-restricted gene expression. We uncovered an interplay between enhancer activity and silencer activity critical for proper regulation of FOXP3 expression dynamics, which could prove generalizable to regulation of other lineage-defining factors.

Overall, this work comprehensively characterized FOXP3 CREs in Tregs and identified multiple novel CREs that enhance or limit FOXP3 expression in human Tconvs. Additionally, we identified TFs and chromatin modifiers that promote and repress FOXP3 expression, and characterized the role of select TFs in maintaining accessibility at FOXP3 enhancers in Tconv.

This work adds new complexity to regulation governing a gene indispensable to immune homeostasis by uncovering novel CREs selectively active in Tconvs and highlighting species-specific differences in regulation at the FOXP3 locus. We expect these insights will inform ongoing efforts to enhance safety, stability, and efficacy of both natural Treg and Tconv-derived Treg cellular immunotherapies for autoimmune disease, transplant tolerance, and graft-versus-host-disease by introducing novel routes to manipulate in situ CREs to enforce or silence FOXP3 expression. The combined power to map CREs and critical TFs that regulate these elements, and to tune these circuits with targeted chromatin modification, provides broad opportunities for cell and gene therapies at diverse loci and cell types.

### Limitations of study

We deciphered regulation of FOXP3 in Tregs and Tconvs in states of rest and activation, but additional cell-states associated with development, TCR engagement, costimulation, cytokine exposure, and distinct in vivo microenvironments may eventually reveal additional unique circuits regulating FOXP3. In addition, despite employing CRE CRISPRi screening at a massive scale (tiling across >120kb in the *FOXP3* locus), our screens may not capture the full region of potential distal CREs that influence FOXP3. Broader cis-screening libraries or selective targeting of additional active chromatin regions may provide additional insight into elements with potential to influence FOXP3.

## RESOURCE AVAILABILITY

### Lead contact

Correspondence and requests for material should be addressed to Alexander Marson.

### Materials availability

This study did not generate new materials.

### Data and code accessibility

Sequencing data is available upon request and will be deposited to NCBI Gene Expression Omnibus prior to publication. Code is available upon request and will be deposited to a public repository prior to publication. PacBio long-read sequencing transcript annotation is available at https://downloads.pacbcloud.com/public/dataset/Kinnex-single-cell-RNA/.

## ACKNOWLEDGMENTS

We thank all members of the Marson Lab for valuable input and discussion. F. Urnov read the manuscript and offered important suggestions. We thank Jane Srivastava of the Gladstone Flow Cytometry Core for technical help and support with FACS, which is supported by NIH S10 RR028962 and the James B. Pendleton Charitable Trust. We thank Dr. Shimon Sakaguchi for providing pre-processed PBAT-seq analysis files. We also acknowledge the Juvenile Diabetes Research Foundation and the Larry L. Hillblom Foundation for their support of this work. A.M. received funding from the Simons Foundation, Lloyd J. Old STAR Award (Cancer Research Institute), Parker Institute for Cancer Immunotherapy, Innovative Genomics Institute, Larry L. Hillblom Foundation (grant no. 2020-D-002-NET), and Northern California JDRF Center of Excellence. A.M. received gifts from Karen Jordan, the Caulfield family, the Byers family, and the CRISPR Cures for Cancer Initiative. S.K. was funded by NIH T32AR050942 and the Lupus Foundation of America Career Development Award. S.K. is an Ernest and Amelia Gallo Endowed Fellow of the Stanford Maternal and Child Health Research Institute. J.A.B. was supported by the Hanna Gray Fellow program of the Howard Hughes Medical Institute. C.T.M. is a UCSF ImmunoX Computational Immunology Fellow, is supported by NIH grant F30AI157167, and has received support from NIH grants T32DK007418 and T32GM007618. B.G.G. was supported by the IGI-AstraZeneca Postdoctoral Fellowship. J.W.F. was funded by US National Institutes of Health (NIH) grant R01HG008140. J.E.C. is supported by the NOMIS Foundation, the Lotte and Adolf Hotz-Sprenger Stiftung, the Swiss National Science Foundation (project grants 310030_188858 and 310030_201160), and the European Research Council (ERC) under the European Union’s Horizon 2020 research and innovation program (grant agreement No 855741, DDREAMM). A.T.S. was supported by a Lloyd J. Old STAR Award from the Cancer Research Institute, the Parker Institute for Cancer Immunotherapy, and the CRISPR Cures for Cancer Initiative.

## AUTHOR CONTRIBUTIONS

J.M.U. and A.M. conceptualized the study. J.M.U, M.M.A., R.D., S.S., and N.K. conducted human cell culture and human experiments. J.M.U. and V.N. sorted cells in screening experiments. D.R.S, B.G.G., G.L.C., and J.E.C designed and generated the CRISPRi library. S.K., W.Z., and R.Z. conducted and analyzed ChIP-seq. A.T.S. supervised ChIP-seq experiments. J.A.B. performed ATAC-seq analysis. H.Y.C. supervised ATAC-seq analysis. J.W.F. developed code for motif analyses. J.M.U., S.S., C.T., R.H., and Z.L. conducted murine experiments. S.W. conducted differential motif analysis. K.P. advised and supervised differential motif analysis. J.M.U. performed general data analysis and visualization. M.M.A. and C.T.M provided analysis direction. L.G. and L.A.G. provided conceptual direction. M.O. provided statistical direction. M.M.A. helped design experiments. J.M.U wrote the original manuscript. J.M.U and A.M. edited the manuscript. A.M. provided resources, acquired funding, and supervised the study.

## METHODS

### Primary human T cell isolation and culture

Primary human T cells subsets were isolated from peripheral blood mononuclear cells (PBMCs) sourced from consented, fresh Human Peripheral Blood Leukopaks (STEMCELL Technologies, catalog no. 70500). Leukopaks were washed twice with 1X volume of EasySep Buffer (DPBS, 2% fetal bovine serum [FBS], 1 mM pH 8.0 EDTA) using centrifugation and resuspended at 50e6-200e6 cells/mL. CD4+CD25^high^CD127^low^ Tregs and CD4+CD25^low^ Tconv were isolated from washed PBMCs using the EasySep Human CD4+CD127lowCD25+ Regulatory T Cell Isolation Kit (STEMCELL Technologies, catalog no. 18063) according to manufacturer’s protocol. For enhanced cell purity, Tregs were stained for CD4 (Biolegend, catalog no. 344634 or 344620), CD25 (Tonbo, catalog no. 20-0259-T100), and CD127 (Becton Dickinson, catalog no. 557938) and further sorted into a CD4+CD25^high^CD127^low^ population via FACS on a BD FACSAria or BD FACSAria Fusion I. Tregs and Tconvs were cultured in X-VIVO15 media (Lonza, catalog no. 02-053Q) supplemented with 5% FBS, 55 µM 2-Mercaptoethanol, and 4 mM N-Acetyl-L-Cysteine. For CRISPRi screening, CRISPRi validation, and CRISPRn screening, recombinant Human IL-2 (R&D Systems, catalog no. 202-GMP or BT-002-GMP) was supplemented at 200 IU/mL, and cells were stimulated for 48 hours with CTS Dynabeads CD3/CD28 beads (Thermo Fisher Scientific, catalog no. 40203D) at a cell to bead ratio of 1:1.

After 48 hours, beads were removed using magnetic separation. For all other experiments, cells were cultured using recombinant Human IL-2 (R&D Systems, catalog no. 202-GMP or BT-002- GMP) supplemented at 300 IU/mL and stimulated with 12.5 uL/mL Immunocult Human CD3/CD28/CD2 T Cell Activator (STEMCELL Technologies, catalog no. 10970), to enhance cell recovery, growth, and viability. Cells were cultured at 37°C and 5% CO_2_ and split to 5E5-1.5E6 cells/mL every ∼48 hours by topping off media and completely replacing the appropriate dose of IL-2.

### Libraries and plasmids

The CRISPRi gRNA library was designed and cloned spanning approximately chrX:49,225,440- 49,348,360 (hg38), as previously described^7^, and contains ∼15K gRNAs flanked by a 5’-NGG protospacer adjacent motifs (Table S1). Briefly, gRNA sequences with cloning adapters were synthesized by Agilent Technologies, amplified, and cloned into pCRISPRia-v2 (Addgene, 84832). The dCas9-ZIM3 plasmid was designed and cloned, as previously described^7^. The CRISPRn gRNA library contains 6000 gRNAs targeting 1349 TFs, chromatin modifiers, and immune genes of interest, with 593 non-targeting controls and 13 EGFP-targeting controls^26^.

### Lentivirus production

Lentiviruses containing the CRISPRi gRNA library, dCas9-ZIM3 construct, and CRISPRn gRNA library were generated as previously described^48^. Low passage number Human Embryonic Kidney 293T cells were thawed and cultured at 37°C and 5% CO_2_ in DMEM, high glucose, GlutaMAX Supplement medium (Fisher Scientific, catalog no. 10566024) supplemented with 10% FBS, 100 U/mL Penicillin-Streptomycin (Fisher Scientific, catalog no. 15140122), 2 mM L-glutamine (Fisher Scientific, catalog no. 25030081), 10 mM HEPES (Sigma, catalog no. H0887-100ML), 1X MEM Non-Essential Amino Acids Solution (Fisher Scientific, catalog no. 11140050), and 1 mM Sodium Pyruvate (Fisher Scientific, catalog no. 11360070) for at least 3 passages, without exceeding 80% confluency. On day 0, 293T cells were seeded in a flat-bottom culture vessel at medium-high confluency in Opti-MEM I Reduced Serum Medium (Fisher Scientific, catalog no. 31985088) supplemented with 5% FBS, 100 U/mL Penicillin-Streptomycin (Fisher Scientific, catalog no. 15140122), 2 mM L-glutamine (Fisher Scientific, catalog no. 25030081), 1X MEM Non-Essential Amino Acids Solution (Fisher Scientific, catalog no. 11140050), and 1 mM Sodium Pyruvate (Fisher Scientific, catalog no. 11360070) to achieve 95% confluency following overnight incubation at 37°C and 5% CO_2_ overnight. When cells reached approximately 95% confluency, transfection complexes were assembled. To assembled transfection complexes, supplement-free Opti-MEM (Fisher Scientific, catalog no. 31985088) was adjusted to room temperature. Lipofectamine3000 Mastermix was assembled by adding 0.79 µL Lipofectamine 3000 reagent (Fisher Scientific, catalog no. L3000075) per cm^2^ of the 293T culture vessel to 1/8 293T culture volume of room temperature supplement-free Opti-MEM. In parallel, P3000 Mastermix was assembled by adding 125 ng psPAX2 (Addgene 12260) per cm^2^ of the 293T culture vessel, 62.5 ng pMD2.G (Addgene 12259) per cm^2^ of the 293T culture vessel, and 167 ng transfer plasmid (CRISPRi gRNA library, dCas9-ZIM3, or CRISPRn gRNA library) per cm^2^ of the 293T culture vessel to 1/8 293T culture volume of room temperature supplement-free Opti-MEM. After addition of plasmids, 0.71 µL p3000 Reagent (Fisher Scientific, catalog no. L3000075) per cm^2^ of the 293T culture vessel was added.

Mastermixes were mixed gently by inversion, and Lipofectamine3000 Mastermix was added dropwise to P3000 Mastermix, gently inverting to mix. The combined transfection mix was incubated at room temperature for 15 min. One-fourth volume of the 293T medium was removed, and the equivalent volume of transfection complex was added to 293T cultures and incubated for 6 hours at 37°C and 5% CO_2_. After incubation, media was removed and replaced with fresh complete media with 1X ViralBoost Reagent (Alstem, catalog no. VB100) and incubated 18 hours at 37°C and 5% CO_2_. Twenty-four hours after transfection, media was transferred to 50 mL centrifuge tubes and centrifuged at 300g for 5 min to remove cell debris. Supernatant was transferred to a new tube and stored at 4°C, and fresh complete media with 1X ViralBoost was gently replaced on 293T. Forty-eight hours post-transfection, media was collected again, as described, and combined with supernatant collected at 24-hours post-transfection. Lenti-X Concentrator (Takara Bio, catalog no. 631232) was added to combined supernatant, and lentiviral particles were concentrated according to manufacturer’s protocol and resuspended in supplement-free Opti-MEM to 1% of the original culture volume. Lentiviral particles were aliquoted and frozen at -80°C, thawing immediately prior to use.

### Intracellular flow cytometry staining

Approximately 5E4-3E5 cells per donor and cell type were transferred to a 96-well V-bottom plate, centrifuged at 300g for 5 min, and supernatant was removed. Cells were washed with 200 µL EasySep buffer and resuspended in 50 µL of staining solution containing Ghost Dye Red 780 Live/Dead stain (Tonbo, catalog no. 13-0865-T500) and antibodies targeting surface proteins of interest. Cells were incubated on ice for 20 min, protected from light. After staining, cells were washed by addition of 150 µL EasySep, centrifuged at 300g for 5 min, and supernatant was removed. Intracellular staining was conducted using the FOXP3 Fix/Perm Buffer Set (Biolegend, catalog no. 421403). Kit components were diluted in DPBS according to manufacturer’s protocol. Cells were resuspended in 50 µL 1X FOXP3 Fix/Perm Buffer and incubated at room temperature for 30 min, protected from light. After fixation, cells were washed by addition of 150 µL 1X FOXP3 Perm Buffer, centrifuged at 300g for 5 min, and supernatant was removed.

Cells were permeabilized in 200 µL 1X FOXP3 Perm Buffer for 15 min at room temperature, protected from light. Following permeabilization, cells were centrifuged at 300g for 5 min, and supernatant was removed. Cells were resuspended in 50 µL 1X FOXP3 Perm Buffer containing antibodies targeting intracellular proteins of interest and incubated 30 min at room temperature, protected from light. Following intracellular staining, cells were washed by addition of 150 µL 1X FOXP3 Perm Buffer, centrifuged at 300g for 5 min, and supernatant was removed. Cells were resuspended in 200 µL EasySep for flow cytometry. Stained cells were analyzed on an Attune NxT Flow Cytometer. For larger samples, volumes were adjusted to accommodate staining of higher cell numbers in 50 mL conical tubes.

### CRISPRi tiling screen

CRISPRi screens were conducted in Tconv and Treg from the same two donors, with a third donor used only in Tconv. Of note, Donor 2 in the Tconv screen showed no low bin enrichment at the *FOXP3* TSS, indicating a failed experiment, and was excluded from the combined donor analysis described above.

*Lentiviral infection and selection.* Twenty-four hours post-stimulation, cells were infected with titered dCas9-ZIM3 lentivirus, targeting approximately 80% infection, by addition to culture flasks. Cells and lentivirus were gently pipetted once with a large serological pipette to mix.

Forty-eight hours post-stimulation, cells were infected with gRNA library lentivirus, targeting approximately 60% infection, by addition to culture flasks and gently mixing. Twenty-four hours post-gRNA library lentivirus infection, viral media was removed, and cells were resuspended in media containing 1.5 ug/mL puromycin to select for cells with proper gRNA integration. Cells were cultured in puromycin-supplemented media for 48 hours.

*Cell purity intracellular staining.* Seven days post-stimulation, a small fraction of Treg and Tconv were removed from culture to assess cell population purity, and stained as described in *Intracellular flow cytometry staining*. Cells were stained using antibodies targeting CD4 (Biolegend, catalog no. 344634), FOXP3 (Biolegend, catalog no. 320112), and HELIOS (Biolegend, catalog no. 137216). Cells were gated on live cells and CD4+, and the percentage of FOXP3+HELIOS+ cells (Tregs) were compared to the percentage of FOXP3-HELIOS- cells (Tconvs).

*Restimulation and cell sorting.* Nine days post-stimulation, cells were restimulated at a 1:1 cell to bead ratio with CTS Dynabeads CD3/CD28 (Thermo Fisher Scientific, catalog no. 40203D).

Forty-eight hours post-restimulation, beads were separated from cells using magnetic isolation. Tregs and Tconvs were counted, washed, and stained with Ghost Dye Red 780 Live/Dead stain (Tonbo, catalog no. 13-0865-T500) and antibodies targeting CD4 (Biolegend, catalog no. 344634), FOXP3 (Biolegend, catalog no. 320112) and HELIOS (Biolegend, catalog no. 137216), as described in *Intracellular flow cytometry staining*, with adjusted volumes to accommodate large cell numbers. Cells were sorted on lymphocytes, singlets, live cells, BFP+ (a marker of the gRNA library plasmid), and FOXP3 high and low bins that captured the top and bottom ∼25% of FOXP3 expression in each cell type. Following sorting, cells were pelleted and lysed, and genomic DNA (gDNA) was extracted using phenol-chloroform gDNA extraction.

*Library preparation and sequencing.* Regions containing the lentivirally integrated gRNA were amplified using PCR with custom primers. PCR reactions were conducted using 2 µg gDNA per 50 µL reaction with 0.5 µM forward and reverse primers, 0.075 U/µL TaKaRa Ex Taq DNA polymerase (Takara Bio, catalog no. RR001C), 0.2 mM dNTP, and 1X Ex Taq Buffer and amplified under the following cycling conditions: 1 min at 95°C, (30 s at 95°C, 30 s at 60°C, 30 s at 72°C) x 25 cycles, 10 min at 72°C, hold at 4°C. Following amplification, aliquots of individual samples were pooled, and primer dimers removed using a 1.4X SPRI bead cleanup.

Proper amplification and concentration were assessed using Qubit quantification and DS1000 High Sensitivity Tapestation analysis, according to manufacturers’ protocols. Samples were pooled equimolarly and sequenced on an Illumina HiSeq4000 instrument using a custom sequencing primer.

### CRISPRi screen analysis

To acquire genomic coordinates of the CRISPRi library gRNAs, gRNA sequences were mapped to the expanded *FOXP3* locus (chrX:49200000-49348394, human genome hg38) using bwa (v0.7.18)^49^ with the following command and specifications: bwa aln -n 0. Aligned SAM files were converted to BAM files and indexed using samtools (v1.10) and converted to a BED file containing genomic coordinates using bedtools (v2.31.1) bamtobed. 14,889 gRNAs were successfully mapped, and the midpoint of each gRNA was selected as the coordinate for plotting.

CRISPRi pooled screens were analyzed using MAGeCK^50^ (v0.5.9.5). A count file containing individual gRNA abundance across all donors was generated using the command mageck count with specification –norm-method none (Table S2). Differential enrichment of gRNAs was determined using DESeq2 (v1.40.2), as described previously^7^ (Table S3, Table S4). Guide RNAs with fewer than ten reads were excluded. Guide RNAs not mapping to the *FOXP3* locus were excluded from visualization.

To decrease noise and enable comparison of individual donor replicates, count files were generated for individual donor samples, as described in combined analyses. Guide RNAs not mapping to the *FOXP3* locus were excluded. Neighboring gRNAs were grouped into sliding 500 bp bins, shifted 50 bp at a time, as “genes”, and gRNAs with less than 10 reads across 5+ samples were excluded from analysis. Guide RNA “gene” bins from individual donors with significant differential enrichment between low and high FOXP3 FACS bins were identified using the binned count file with the command mageck test.

### Cas9 RNP assembly

To assemble RNPs for individual edits and paired deletions, individual custom crRNAs targeting regions of interest (Dharmacon) and Edit-R CRISPR-Cas9 Synthetic tracrRNA (Dharmacon, catalog no. U-002005-20) were resuspended in Nuclease-Free Duplex Buffer (IDT, catalog no. 11-01-03-01) at 160 uM, complexed at a 1:1 molar ratio to create a 80 µM solution of RNA complexes, and incubated 30 min at 37°C. Single-stranded oligonucleotides (ssODN; 100 µM stock, sequence: TTAGCTCTGTTTACGTCCCAGCGGGCATGAGAGTAACAAGAGGGTGTGGTAATATTACGGTACCGAGCACTATCGATACAATATGTGTCATACGGACACG) were mixed at a 1:1 molar ratio with complexed RNA and incubated 5 min at 37°C. Cas9 protein (Berkeley Macrolab, 40 µM stock) was slowly added at a 1:1 molar ratio of Cas9 to RNA complexes, mixed thoroughly, and incubated 15 min at 37°C. For RNPs used in paired deletions, individually assembled RNPs were mixed 1:1 with the appropriate RNP pair. RNPs were frozen at -80°C and thawed at room temperature prior to use.

### RNP electroporation

Cells (Tregs and/or Tconvs) were isolated and stimulated as described above. Forty-eight hours post-stimulation, cells were counted, pelleted via centrifugation, and resuspended at 2E4-7.5E4 cells/µL in freshly supplemented P3 Primary Cell Nucleofector Solution (Lonza, catalog no. V4SP-3096). For single RNP edits, 20 µL cells were mixed with 3.5 µL RNP, and 22 µL was transferred to a 96-well Nucleocuvette Plate (Lonza, catalog no. V4SP-3096). For paired RNP deletions, 20 µL cells were mixed with 7 µL paired RNPs, and 24-24.5 µL was transferred to a 96-well Nucleocuvette Plate (Lonza, catalog no. V4SP-3096). Cells were nucleofected using a Lonza 4D 96-well electroporation system with pulse code EO-115 (human Treg), EH-115 (human Tconv), or CM-137 (murine Tconv). Immediately after electroporation, 80 µL pre-warmed media was added to each well, and the cells were incubated for 15 min at 37°C. Following incubation, cells were plated at approximately 1E6 cells/mL for culture.

### CRISPRi screen validation

*Cas9 RNP assembly and electroporation.* To validate CRISPRi screening in an arrayed format, paired crRNAs tiling CRISPRi-responsive and control regions were designed to delete approximately 1 kb of DNA. Single (3) and paired (2) crRNAs targeting the *AAVS1* safe-harbor locus were designed as negative controls, and single crRNAs from the Brunello Library targeting *FOXP3* exons (4) were included as positive controls^51^. Additional single crRNAs from the Brunello Library targeting *PPP1R3F* exons (4) were included to control for effects of gene KO when tiling across the TSS region^51^. RNPs were assembled as described in *Cas9 RNP assembly*. Forty-eight hours post-stimulation, Tregs and Tconv were electroporated as described in *RNP electroporation*.

*Flow cytometry analysis of arrayed validation.* Nine days post-stimulation, a portion of cells were stained with Ghost Dye Red 780 Live/Dead stain (Tonbo, catalog no. 13-0865-T500) and antibodies targeting CD4 (Biolegend, catalog no. 344620), CD25 (Tonbo, catalog no. 20-0259- T100), FOXP3 (Biolegend, catalog no. 320112), and HELIOS (Biolegend, catalog no. 137216), as described in *Intracellular flow cytometry staining*, above. Remaining unstained cells were restimulated with CTS Dynabeads CD3/CD28 beads (Thermo Fisher Scientific, catalog no.

40203D) at a cell to bead ratio of 1:1. Forty-eight hours post-restimulation, beads were removed via magnetic separation, and cells were stained as described on day 9. Flow cytometry data from stained cells was collected on an Attune NxT Flow Cytometer. Analysis with gating on lymphocytes, singlets, live cells, and CD4+ cells was conducted using FlowJo. Flow cytometry statistics were exported via the FlowJo Table Editor and visualized in R using ggplot2 (v3.5.1). For arrayed validation dot plots, log2 of FOXP3 MFI (Tregs) or %FOXP3+ cells (Tconvs) in KOs from each donor over the mean of the donor-matched *AAVS1* controls was shown.

### Genotyping of arrayed paired gRNA deletions

Genomic DNA from 3E4-2E5 cells was isolated from single edit and paired deletion samples using QuickExtract DNA Extraction Solution (Biosearch Technologies, catalog no. QE09050), using manufacturer’s protocol with reduced reagent volume to accommodate processing in a 96- well PCR plate. Paired PCR primers were designed approximately 200 bp upstream and downstream of the deleted region cut sites, with some regions requiring smaller or larger distances due to locus-specific sequence limitations. Paired primers were used to amplify the deleted region in gDNA from deletion samples and control (unedited) gDNA. PCR reactions were conducted using 3 uL QuickExtract-isolated gDNA per 50 uL reaction, 0.5 uM forward and reverse primers, and 1X NEBNext Ultra II Q5 Master Mix (New England Biolabs, catalog no. M0544). Reactions were amplified under the following cycling conditions: 3 min at 98°C, (10 s at 98°C, 20 s at 69°C, 1 min at 72°C) x 31 cycles, 5 min at 72°C, hold at 4°C. Following amplification, primer dimers were removed using SPRI bead cleanup, and the proportion of bands with versus without deletion was approximately quantified using Tapestation analysis, according to manufacturers’ protocols, and compared to control (unedited) gDNA. Bands were assigned as “deletion” or “no deletion” bands if the reported peak size was the expected band size +/- 20%. Bands outside this range were excluded, and the percent moles were calculated based on ratios of “deletion” and “no deletion” bands only. Genotyping ratios are approximate due to PCR bias of differing band sizes. Editing efficiency of individual gRNA edits was assessed using ICE analysis^52^, and the KO-Score is reported.

### RNA-seq visualization

Pre-processed RNA-seq deduplicated counts^53^ from resting and stimulated Tregs and Tconvs were downloaded from GEO (GSE276096, GSE276096_dedup_counts_all.csv), subset to *AAVS1* control samples, and normalized using DESeq2 (v1.40.2), as previously described^53^. Normalized counts of FOXP3, FLICR, and PPP1R3F from *AAVS1*-targeted control Tregs and Tconvs were plotted in R using ggplot2 (v3.5.1).

### ATAC-seq analysis

Fastqs from previously published ATAC-seq datasets^22,26^ corresponding to unstimulated and stimulated CD4 Effector T cells (Tconv) and Regulatory T cells (Treg)^22^ or KO ATAC-seq datasets in CD4+CD25- (Tconv)^26^ were downloaded from GEO series GSE118189 and GSE171737, respectively. Fastq files were processed as previously described^54^. Briefly, adapters were trimmed using fastp, and reads were aligned to the hg38 reference genome using hisat2.

Reads were deduplicated using picard. Peaks were called from each sample using MACS2. ATAC fragments for each sample were converted into bigwig files normalized by the number of reads in transcription start sites in each sample using ‘rtracklayer::export’ in R. Normalized bigwigs from a representative donor (GEO Series GSE118189: GSM3320328 [1003- Effector_CD4pos_T-U], GSM3320329 [1003-Effector_CD4pos_T-]), GSM3320338 [1003- Regulatory_T-U], GSM3320339 [1003-Regulatory_T-S]; GEO Series GSE171737: GSM5232967 [Donor_4_AAVS1_3_H2_ATAC], GSM5232978 [Donor_4_GATA3_F9_ATAC], GSM5232992 [Donor_4_STAT5A_G7_ATAC], GSM5232993, [Donor_4_STAT5B_G2_ATAC], GSM5232980 [Donor_4_IL2RA_H1_ATAC], GSM5232984 [Donor_4_JAK3_H3_ATAC]) were visualized in R using ggplot2 (v3.5.1) using a sliding binning strategy with a bin size of 100 and step size of 20. Differential accessibility was determined using DESeq2 (v1.40.2) with significance cutoffs of *Padj* ≤ 0.1 and *Padj* ≤ 0.05 (Table S10).

### PBAT-seq visualization

Pre-processed PBAT-seq files (from Figure 1E of ref^24^) were provided for visualization at the *FOXP3* locus. PBAT-seq tracks from representative donor STR1 were visualized using a sliding binning strategy with a bin size of 1500 and step size of 300 in ggplot2 (v3.5.1). The following cell populations described in ref^24^ are labeled as follows: Fr1, Naïve Treg; Fr2, Activated Treg; Fr5, Activated Tconv, Fr6, Naïve Tconv.

### CRISPRn screen

CRISPRn screens were conducted in Tregs from two healthy donors and Tconvs isolated from three healthy donors. Screens were conducted as previously described^26^, with minor modifications.

*Lentiviral infection and electroporation.* Cells were isolated and stimulated, as described above. Twenty-four hours post-stimulation, cells were infected with titered gRNA library lentivirus, targeting approximately 85% infection, by addition to culture flasks. Cells and lentivirus were gently pipetted once with a large serological pipette to mix. Twenty-four hours post-gRNA library lentivirus infection, viral media was removed, and cells were washed with 1X volume pre-warmed media and plated at approximately 1E6 cells/mL. Cas9 RNPs were assembled as described in *Cas9 RNP assembly*, above, using a Guide Swap crRNA method^55^ (Edit-R crRNA nontargeting Control 3, Dharmacon, catalog no. U-007503-01-05). Seventy-two hours post-stimulation, cells were pelleted, and resuspended at 5E4 cells/µL (Treg) or 7.5E4 cells/µL (Tconv) in freshly supplemented P3 Primary Cell Nucleofector Solution (Lonza, catalog no.

V4SP-3096). 20 µL of cell were mixed with 3.5 µL (50 pmol; Tregs) or 7 µL (100 pmol; Tconvs) RNP, and 22 µL (Tregs) or 25 µL (Tconvs) was transferred to a 96-well Nucleocuvette Plate (Lonza, catalog no. V4SP-3096) and electroporated, as described in *RNP electroporation*, above.

*Restimulation and cell sorting.* Nine days post-stimulation, cells were restimulated at a 1:1 cell to bead ratio with CTS Dynabeads CD3/CD28 (Thermo Fisher Scientific, catalog no. 40203D).

Forty-eight hours post-restimulation, beads were separated from cells using magnetic isolation. Cells were counted and stained with Ghost Dye Red 780 Live/Dead stain (Tonbo, catalog no. 13- 0865-T500) and antibodies targeting FOXP3 (Biolegend, catalog no. 320116) and HELIOS (Biolegend, catalog no. 137216), as described in *Intracellular flow cytometry staining*, with adjusted volumes to accommodate large cell numbers. Cells were sorted on lymphocytes, singlets (x2), live cells, GFP+ (a marker of the gRNA library plasmid), and FOXP3 high and low bins that captured the top and bottom 25% of FOXP3 expression. Following sorting, cells were pelleted and lysed, and gDNA was extracted using phenol-chloroform gDNA extraction.

*Library preparation and sequencing.* Regions containing the lentivirally integrated gRNA were amplified using PCR with custom primers. PCR reactions were conducted using 1.75 µg gDNA per 50 µL reaction with 0.25 µM forward and reverse primers and 1X NEBNext Ultra II Q5 Master Mix (New England Biolabs, catalog no. M0544). Reactions were amplified under the following cycling conditions: 3 min at 98°C, (10 s at 98°C, 10 s at 63°C, 25 s at 72°C) x 23 cycles, 2 min at 72°C, hold at 4°C. Following amplification, aliquots of individual samples were pooled, and primer dimers removed using a 1.25X SPRI bead cleanup. Proper amplification and concentration were assessed using Qubit quantification and DS1000 High Sensitivity Tapestation analysis, according to manufacturers’ protocols. Samples were pooled equimolarly and sequenced on an Illumina HiSeq4000 instrument using a custom sequencing primer.

### CRISPRn screen analysis

CRISPRn pooled screens were analyzed using MAGeCK^50^ (v0.5.9.5). A count file containing individual gRNA abundance across all donors was generated using the command mageck count with specification –norm-method none (Table S5). Differentially enriched gRNAs and significantly enriched genes in high and low bins were identified using the command mageck test, specifying –sort-criteria pos (Table S6, Table S7, Table S8, Table S9).

### Comparison of murine and human trans-regulator screens

Mageck gene-level output files were downloaded from ref^29^. Mouse and human gene homologies were downloaded from the Mouse Genome Informatics MGI Data and Statistical Reports (ref.^56^) and used to convert mouse gene IDs to human homologs. Two hundred twenty-two genes were successfully converted and overlapped between the libraries. Screen correlation was visualized in R using ggplot2 (v3.5.1).

### CRISPRn screen validation

Top FOXP3 positive and negative regulators were selected for arrayed validation, and two crRNAs were selected for each regulator. Six crRNAs targeting the *AAVS1* locus were additionally included as negative controls. Cas9 RNPs were assembled as described in *Cas9 RNP assembly*, above. Human Tregs and Tconvs from two healthy donors were isolated and stimulated as described above. Forty-eight hours post-stimulation, cells were electroporated with trans-regulator targeting RNPs, as described in *RNP electroporation*, above. Nine days post-stimulation, a portion of cells were stained with Ghost Dye Red 780 Live/Dead stain (Tonbo, catalog no. 13-0865-T500) and antibodies targeting CD4 (Biolegend, catalog no. 344634), FOXP3 (Biolegend, catalog no. 320112), and HELIOS (Biolegend, catalog no. 137216), as described in *Intracellular flow cytometry staining*, above. Remaining unstained cells were restimulated with 12.5 uL/mL Immunocult Human CD3/CD28/CD2 T Cell Activator (STEMCELL Technologies, catalog no. 10970). Forty-eight hours post-restimulation, cells were stained as described on day 9. Flow cytometry data from stained cells was collected on an Attune NxT Flow Cytometer. Analysis with gating on lymphocytes, singlets, live cells, and CD4+ cells was conducted using FlowJo. Flow cytometry statistics were exported via the FlowJo Table Editor and visualized in R using ggplot2 (v3.5.1). For arrayed validation bar plots, raw %FOXP3+ cells or FOXP3 MFI normalized to the mean of the donor-matched *AAVS1* controls were shown.

### Genotyping of arrayed CRISPRn knock-outs

Genomic DNA from 3E4-2E5 cells was isolated from arrayed samples using QuickExtract DNA Extraction Solution (Biosearch Technologies, catalog no. QE09050), using manufacturer’s protocol with reduced reagent volume to accommodate processing in a 96-well PCR plate. Paired PCR primers were designed flanking the gRNA cut site to generate an amplicon of approximately 200 bp. PCR reactions were conducted for each sample using 4 µL QuickExtract- isolated gDNA per 25 µL reaction, 0.5 µM forward and reverse primers, and 1X NEBNext Ultra II Q5 Master Mix (New England Biolabs, catalog no. M0544). Reactions were amplified under the following cycling conditions: 3 min at 98°C, (20 s at 94°C, 20 s at 65-57.5°C (with 0.5°C incremental decreases per cycle), 1 min at 72°C) x 15 cycles, (20 s at 94°C, 20 s at 58°C, 1 min at 72°C) x 20 cycles, 10 min at 72°C, hold at 4°C. Amplified DNA was diluted 1:200, and 1 µL was used per 25 µL of a second PCR amplification with 1 µM forward and reverse primers and 1X NEBNext Ultra II Q5 Master Mix to attached sequencing adapters and indices. Reactions were amplified under the following cycling conditions: 30 s at 98°C, (10 s at 98°C, 30 s at 60°C, 30 s at 72°C) x 12 cycles, 2 min at 72°C, hold at 4°C. Following amplification, samples were pooled in equal volume, and primer dimers were removed using a 1.3X SPRI bead cleanup.

Proper amplification and concentration were assessed using Qubit quantification and DS1000 High Sensitivity Tapestation analysis, according to manufacturers’ protocols. The pooled samples were sequenced on an Illumina MiSeq instrument with PE 150 reads. Analysis of editing efficiency was performed using CRISPResso2^57^ (v2.2.12), using the command CRISPRessoBatch –batch_settings [crispresso_samplesheet] –skip_failed –n_processes 4 – exclude_bp_from_left 0 –exclude_bp_from_right 0 –plot_window_size 10.

### TF binding site analysis

Motif analysis was conducted as described^26^, with minor modifications. Briefly, datasets containing human or human-predicted position weight matrices for human TFs were acquired from JASPAR2020^58^ (v0.99.10) using TFBSTools (v1.38.0) in R or sourced from CisBP (*Homo sapiens*, downloaded 09-29-2020)^59^. Motif sites in a region of interest were identified using ‘matchMotifs’ from motifatchr (v1.22.0) using default settings and specifying the genome as Bsgenome.Hsapien.UCSC.hg38 (v1.4.5).

### ChIP-sequencing

Up to 2.5E7 Tregs were cross-linked in PBS with 1% methanol-free formaldehyde (Thermo, catalog no. 28908) for 10 min at room temperature followed by quenching in glycine at 125 mM final concentration. Cells for YBX1 ChIP-seq were cross-linked in 2 mM disuccinimidyl glutarate (Thermo, catalog no. 20593) in PBS for 30 min at room temperature prior to formaldehyde treatment. Cross-linked cell pellets were snap-frozen in liquid nitrogen and stored at –80°C. Nuclei were isolated from thawed, cross-linked cells via sequential lysis in LB1 (50 mM HEPES-KOH pH 7.5, 140 mM NaCl, 1 mM EDTA, 10% glycerol, 0.5% IGEPAL CA-360, and 0.25% Triton X-100), LB2 (10 mM Tris-HCl pH 8, 200 mM NaCl, 1 mM EDTA, and 0.5 mM EGTA), and LB3 (10 mM Tris-HCl pH 8, 100 mM NaCl, 1 mM EDTA, 0.5 mM EGTA, 1% sodium deoxycholate [NaDOC], and 0.5% N-laurylsarcosine) supplemented with 0.5 mM phenylmethylsulfonyl fluoride (PMSF, Sigma, catalog no. P7626) and 0.5X protease inhibitor cocktail (PIC, Sigma, catalog no. P8340). Chromatin was sheared on a Covaris E220 focused ultrasonicator using 1 mL milliTubes (Covaris, catalog no. 520128) with 140W peak incident power, 5% duty factor, 200 cycles per burst, 6°C temperature setpoint (minimum 3°C, maximum 9°C), fill level 10, and time 12-14 min to obtain target size 200-700 bp. Triton X-100 was added to final concentration of 1% prior to immunoprecipitation for 16 h at 4°C with 2-8 µg of indicated antibodies (Table S11) bound to a 1:1 mixture of protein A and protein G magnetic beads (Thermo, catalog nos. 10001D and 10003D). Bead-bound antibody-chromatin complexes were sequentially washed three times with Wash Buffer 1 (20 mM Tris pH 8, 150 mM NaCl, 1 mM EDTA, 0.5 mM EGTA, 1% Triton X-100, 0.1% sodium dodecyl sulfate [SDS], and 0.1% NaDOC), twice with Wash Buffer 2 (20 mM Tris-HCl pH 8, 500 mM NaCl, 1 mM EDTA, 0.5 mM EGTA, 1% Triton X-100, 0.1% SDS, and 0.1% NaDOC), twice with Wash Buffer 3 (20 mM Tris-HCl pH 8, 250 mM LiCl, 1 mM EDTA, 0.5% IGEPAL CA-360, and 0.5% NaDOC), twice with TET (10 mM Tris-HCl pH 8, 1 mM EDTA, 0.2% Tween-20), and once with TE0.1 (10 mM Tris-HCl pH 8, 0.1 mM EDTA, 0.5 mM PMSF, and 0.5X PIC) supplemented with 0.5 mM PMSF and 0.5X PIC. Beads were resuspended in TT (10 mM Tris-HCl pH 8, 0.05% Tween- 20) prior to on-bead library preparation using the NEBNext Ultra II DNA Library Prep Kit (NEB, catalog no. E7370L) as described previously^60^. ChIP-seq libraries were multiplexed for paired-end sequencing on an Illumina NextSeq 2000 or NovaSeq X instrument.

### ChIP-seq analysis

*ChIP-seq for new targets*. Reads were trimmed to remove adapters and low-quality sequences and aligned to the hg38 reference genome assembly with bwa^49^ (v0.7.17-r1188) before filtering to remove duplicates and low-quality alignments including to problematic genomic regions^61^ using the nf-core/chipseq pipeline^62^ (v2.0.0, doi: 10.5281/zenodo.3240506) with default parameters. Bigwig files were visualized in R using ggplot2 (v3.5.1) and auto scaled to the locus shown using a sliding binning strategy with a bin size of 100 and step size of 20, unless otherwise noted.

*ChIP-seq from public datasets*. Pre-processed ChIP-seq bigwigs were downloaded from ChIP-Atlas^63^. ChIP-Atlas processing uses Bowtie2 for alignment to the human genome GRCh38, SAMtools to convert to BAM format, sort, and remove PCR duplicates, bedtools to calculate coverage scores in reads per million mapped reads, MACS2 to call peaks, and the UCSC BedGraphToBigWig tool to generate bigwig coverage files. Human ChIP-seq data for the indicated cell type were generated in the following papers and are available at the listed GEO sample accession code: BCL11B TALL (primary T-ALL with enhanced BCL11B expression, GSM5024541, ref.^33^), ETS1 Jurkat (Jurkat clone E6-1, GSM449525, ref.^34^), GABPA Jurkat (Jurkat, GSM1193662, ref.^35^), GATA3 Th1 (in vitro differentiated primary human Th1 cells, GSM776558, ref.^37^), HIC1 iTreg (differentiated iTregs from CD4+CD25- cord blood T cells, GSM2663736, ref.^42^), MBD2 K562 (K-562, GSM2527611, ref.^36^), STAT5 Naïve Treg (CD4+CD25+CD45RA+ expanded naïve Tregs, GSM1056923, ref.^43^), STAT5B T cell + IL-2 (human CD4+ T cells stimulated with IL-2 for 1 hour, GSM671402, ref.^38^), TFDP1 MM-1S (MM.1S, GSM2132551, ref.^39^), YY1 DND41 (T-ALL cell line DND-41 treated with DMSO, GSM5282087, ref.^40^), and ZNF143 CUTLL1 (CUTLL1, GSM732907, ref.^41^). Mouse ChIP-seq data for the indicated cell type were generated in the following papers and are available at the listed GEO sample accession code: H3K27ac nCD4+ T cells (naïve CD4+ T cells, GSM6019885, [WT], ref.^64^), H3K27ac Treg (Tregs, GSM6019890, [WT], ref.^64^), GATA3 nCD4+ T cells (naïve CD4+ T cells, GSM523223, strain C57BL/6, ref.^65^), GATA3 naïve Treg (naïve Tregs, GSM523230, strain C57BL/6, ref.^65^), STAT5 CD4+ T cells (CD4+CD44-CD25- T cells treated with IL-2, GSM6283233, strain C57BL/6 [WT], ref^66^), STAT5 Treg (Tregs, GSM5574817, strain Foxp3-GFP [WT], ref^67^), EGR2 CD4+ T cells (CD4+ T cells treated with immobilized anti-CD3 mAb, GSM1126961, strain C57BL/6, ref.^68^). Additional human ChIP-seq datasets from the Epigenome Roadmap Project were acquired with the following SRX IDs: SRX088919 (H3K27ac in naïve CD4+ T cells, ref.^1^), SRX088820 (H3K27ac in stimulated CD4+ T cells, ref.^1^). Bigwig files were visualized in R using ggplot2 (v3.5.1) and auto scaled to the locus shown using a sliding binning strategy with a bin size of 100 and step size of 20, with the exception of Figure 5C EGR2 ChIP-seq (bin size of 25 and step size of 5).

### Cactus alignment & motif differential affinity analysis

Human (hg38), murine (mm10), and select species sequences were extracted (cactus-hal2maf) from the Zoonomia 241-way Cactus hierarchical alignment^69^ with the murine or human genome as reference. The resulting Multiple Alignment Format (MAF) was subset to regions of interest using mafsInRegion (KentUtils) and converted to FASTAs using msa_view (phast^70^). Finally, these FASTAs were converted to VCFs using a custom script. motifbreakR^71^ tested the potential of each variant to disrupt binding affinity of any motif in HOCOMOCO v11^72^. Murine sequence-level multi-species comparisons at TcNS- were generated using MAF files, which were imported in R using ggmsa^73^ (v1.6.0) and visualized using ggplot2 (v3.5.1), converting all nucleotides to uppercase notation.

### Murine Tconv isolation and culture

Spleens and lymph nodes were harvested from Foxp3-Thy1.1 mice and mashed through a 70 µM filter in EasySep, and flow through cells were centrifuged at 300g for 5 min. Supernatant was aspirated, and cells were resuspended at 100E6 cells/mL. CD4+ T cells were isolated from washed splenocytes and lymph nodes cells using the EasySep Mouse CD4+ T Cell Isolation Kit (STEMCELL Technologies, catalog no. 19852), according to the manufacturer’s protocol.

Murine Tregs were depleted from CD4+ T cells using the EasySep Mouse CD90.1 Positive Selection Kit (STEMCELL Technologies, catalog no. 18958). Tconvs were cultured in Kool Aid Complete Media (DMEM High Glucose, Custom “Kool Aid” media, catalog no. UCSFMP-RM- 4591, obtained from UCSF Media Center) supplemented with 10% FBS (R&D Systems, catalog no. S11550), 100 µM 2-Mercaptoethanol (Sigma, catalog no. 63689-25ML-F), 10 mM HEPES (Sigma, catalog no. H0887-100ML), 1 mM Sodium Pyruvate (Fisher Scientific, catalog no.

11360070), 100 IU/mL Penicillin-Streptomycin (Sigma, catalog no. P4333-100ML), 2 mM L-Glutamine (Fisher Scientific, catalog no. 25030081) and 1X MEM (Fisher Scientific, catalog no. 11140050). Cells were cultured in NeutrAvidin (Fisher Scientific, catalog no.31,000) coated (5ug/mL in PBS) 24 well plates. Cells were stimulated with Biotin Hamster Anti-Mouse CD3e clone 145-2C1 (Becton Dickinson, catalog no. 553060) at a concentration of 0.25 ug/mL and Biotin CD28 clone 37.51 (Becton Dickinson, catalog no.553296) at a concentration of 1 mg/mL. Recombinant Human IL-2 (R&D Systems, catalog no. 202-GMP or BT-002-GMP) was supplemented at 50 IU/mL for Tconv cells. Additionally, recombinant Human IL-7 (Fisher Scientific, catalog no. 207-IL-050) at a concentration of 2.5 ng/mL and unconjugated Human IL- 15 (Fisher Scientific, catalog no. 247-ILB-025) at a concentration of 50 ng/mL were added to enhance cell recovery, growth, and viability. Cells were cultured at 37°C and 5% CO_2_ and split to 1E6 cells/mL every ∼2 days by topping off media and completely replacing the appropriate dose of IL-2, IL-7 and IL-15.

### Murine FOXP3 intracellular flow cytometry staining

Approximately 5E4-3E5 cells per donor and cell type were transferred to a 96-well U-bottom plate, centrifuged at 300g for 5 min, and supernatant was removed. Cells were washed with 150 µL EasySep buffer and resuspended in 25 µL of Mouse BD Fc Block (Becton Dickinson, catalog no. 553142) at a 1:50 dilution in EasySep buffer. Upon staining with Fc Block mix for 1 min at 4°C, cells were stained with staining solution containing Ghost Dye Red 780 Live/Dead stain (Tonbo, catalog no. 13-0865-T500) and Thy1.1-APC monoclonal antibody (Fisher Scientific, catalog no. 17-0900-82). Cells were incubated on ice for 20 min, protected from light. After staining, cells were washed by addition of 150 µL EasySep, centrifuged at 300g for 5 min, and the supernatant was removed. Intracellular staining was conducted using the Foxp3/Transcription Factor Staining Buffer Set (Fisher Scientific, catalog no. 00-5523-00). Kit components were diluted in diluent and diH20 according to the manufacturer’s protocol. Cells were resuspended in 100 µL 1X Fixation/Permeabilization buffer and incubated at 4°C for 30 min, protected from light. After fixation, cells were washed by addition of 150 µL 1X Permeabilization Buffer, centrifuged at 300g for 5 min, and the supernatant was removed. Cells were resuspended in 50 µL 1X Permeabilization Buffer containing FOXP3 Monoclonal Antibody (FJK-16s) eFluor 450 (Fisher Scientific, catalog no. 48-5773-82) and incubated for 30 min at 4°C, protected from light. Following intracellular staining, cells were washed by addition of 150 µL 1X FOXP3 Perm Buffer, centrifuged at 300g for 5 min, and supernatant was removed. Cells were resuspended in 200 µL EasySep for flow cytometry. Stained cells were analyzed on an Attune NxT Flow Cytometer.

### Plots and Genomic Tracks

Gene tracks were plotted from a subset of GENCODE hg38 knownGene annotations (human) and GENCODE VM23 mm10 knownGene annotations (mouse) downloaded from the UCSC Genome Browser (ref.^74^) Table Browser and visualized in R using ggplot2 (v3.5.1). Hg38 PhyloP 100way conservation (human) and GRCm38/mm10 PhyloP 60way All conservation (mouse) track data were downloaded from the UCSC Genome Browser Table Browser and visualized in R with ggplot2 using 25 bp window bins, with the exception of conservation in Figure 5B-C and Figure S9A (5 bp window bins used). PacBio long-read sequencing transcript annotations in gff format were downloaded from https://downloads.pacbcloud.com/public/dataset/Kinnex-single-cell-RNA/, and Revio PBMC 2 transcripts were visualized in R using ggplot2 (v3.5.1). EGR2 motif sequence logos were downloaded from JASPAR^75^. Select schematics incorporated images from https://www.biorender.com and NIH BIOART.

### Supplementary Tables

*S1.* CRISPRi gRNA library.

*S2.* CRISPRi screen gRNA counts generated with MAGeCK.

*S3.* CRISPRi screen Treg results generated with DESeq2.

*S4.* CRISPRi screen Tconv results generated with DESeq2.

*S5.* CRISPRn screen gRNA counts generated with MAGeCK.

*S6.* CRISPRn screen Treg gene-level results generated with MAGeCK. *S7.* CRISPRn screen Treg gRNA-level results generated with MAGeCK. *S8.* CRISPRn screen Tconv gene-level results generated with MAGeCK. *S9.* CRISPRn screen Tconv gRNA-level results generated with MAGeCK.

*S10.* KO ATAC-seq differentially accessible peaks generated with DESeq2.

*S11*. List of ChIP-seq antibodies.

*S12.* List of gRNAs used in arrayed experiments.

---

**Figure S1.**
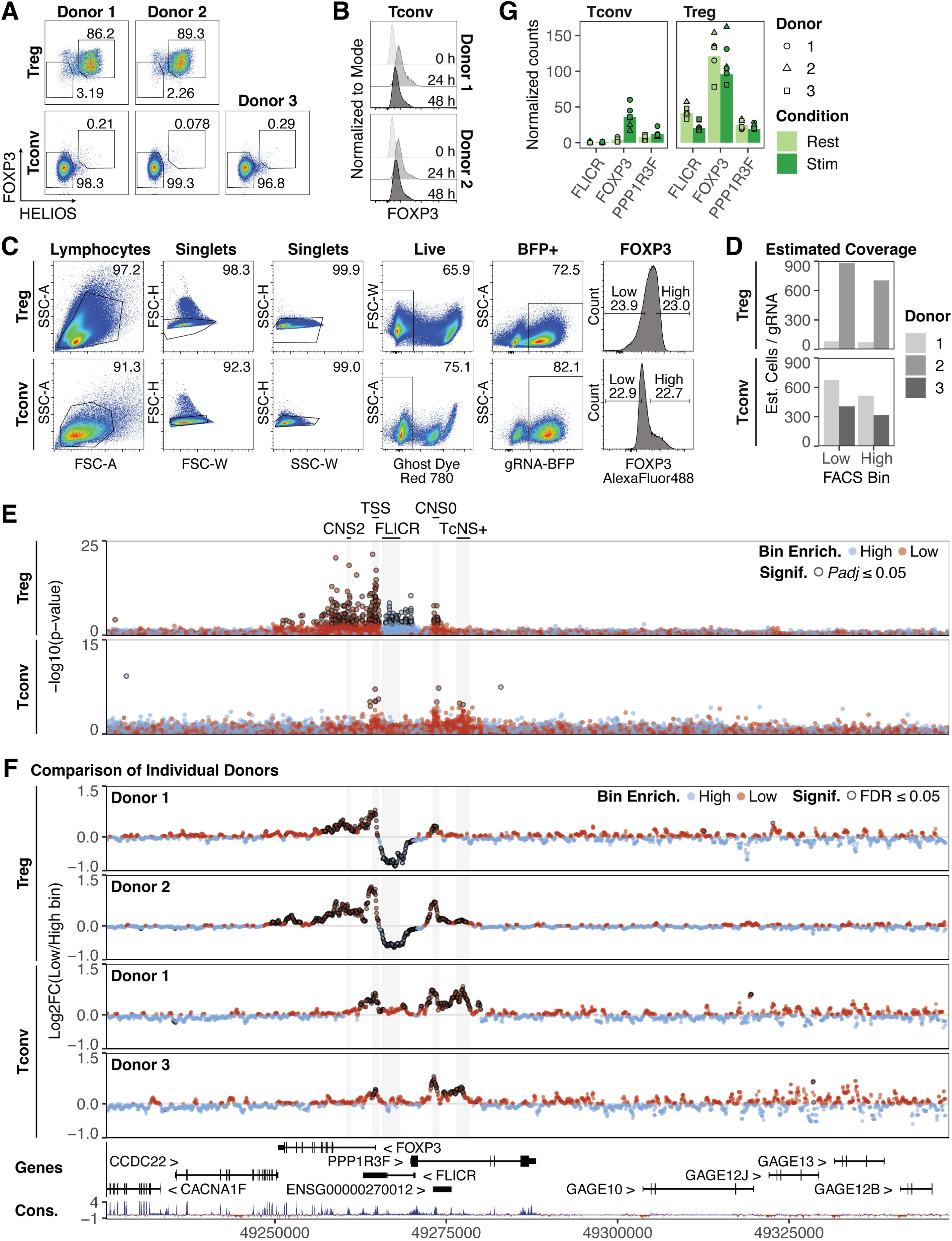
CRISPRi *FOXP3* locus tiling screen identifies FOXP3 CREs. (A) Flow cytometry plots depicting FOXP3 and HELIOS expression in CRISPRi tiling screen Treg and Tconv donors 7 days post-initial stimulation. (B) Flow cytometry plots of FOXP3 expression in *AAVS1*-targeted Tconvs at 0, 24, and 48 hours post-restimulation with anti-CD28/CD3/CD2 antibody complexes. (C) Example representative gating strategy of Treg (top) and Tconv (bottom) FOXP3 expression in CRISPRi tiling screen. (D) Estimated cell coverage per gRNA in FOXP3 high and low bins in Tregs (top) and Tconvs (bottom). Estimated coverage was calculated as the number of sorted cells in each bin divided by 15,029, the theoretical number of gRNAs in the FOXP3 tiling library. (E) CRISPRi *FOXP3* locus-tiling -log10(p-value) of gRNA enrichment in FOXP3 high vs. low FACS bins in Treg and Tconv. Blue, gRNA enriched in FOXP3 High FACS bin. Red, gRNA enriched in FOXP3 Low FACS bin. Outlined, adjusted p-value ≤ 0.05. Treg, *n* = 2 donors; Tconv, *n* = 2 donors. (F) CRISPRi *FOXP3* locus-tiling screen log2 fold change gRNA enrichment in FOXP3 low vs. high FACS bins in individual Treg and Tconv donors, plotted along the *FOXP3* locus. Neighboring gRNAs were grouped into sliding 500 bp bins, shifted 50 bp at a time, for analyses, and individual points indicate grouped bins. Blue, gRNA bin enriched in FOXP3 High FACS bin. Red, gRNA bin enriched in FOXP3 Low FACS bin. Outlined, FDR ≤ 0.05. (G) RNA-seq^53^ normalized counts of FLICR, FOXP3, and PPP1R3F RNA expression in resting and stimulated Tregs and Tconvs.

**Figure S2.**
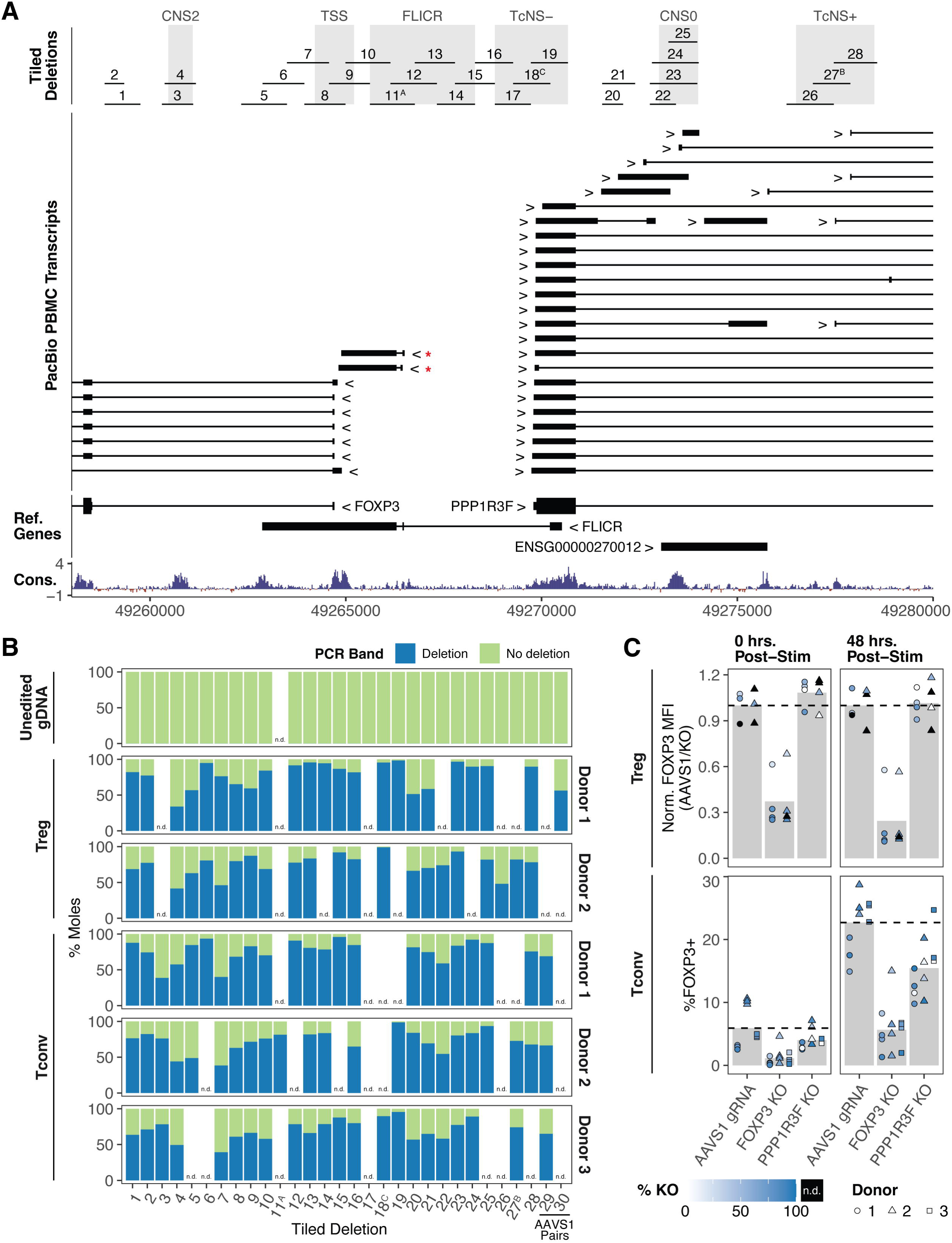
Location and efficiency of CRISPRn paired deletions across the *FOXP3* locus. (A) Top, location of tiled deletions at the FOXP3 locus. Superscript letters correspond to tile labels in the main text and Figure 1. Bottom, transcripts from publicly available long-read PacBio sequencing in human PBMCs. Red stars indicate shortened FLICR isoforms relative to the reference genes. (B) Approximate molar ratio of deletion to no deletion PCR bands in PCR genotyping assays for tiled deletion efficiency. Numbers correspond to Tiled Deletions shown in (A). n.d., no data. (C) Normalized FOXP3 MFI in Tregs (top) and percentage FOXP3+ Tconv (bottom) with *AAVS1* gRNA, PPP1R3F KO, or FOXP3 KO. Point color indicated editing efficiency of individual gRNA (Tconv, *n* = 3 donors; Treg, *n* = 2 donors, x 4 gRNAs per target or 3 *AAVS1*- targeting gRNA controls; n.d., no data).

**Figure S3.**
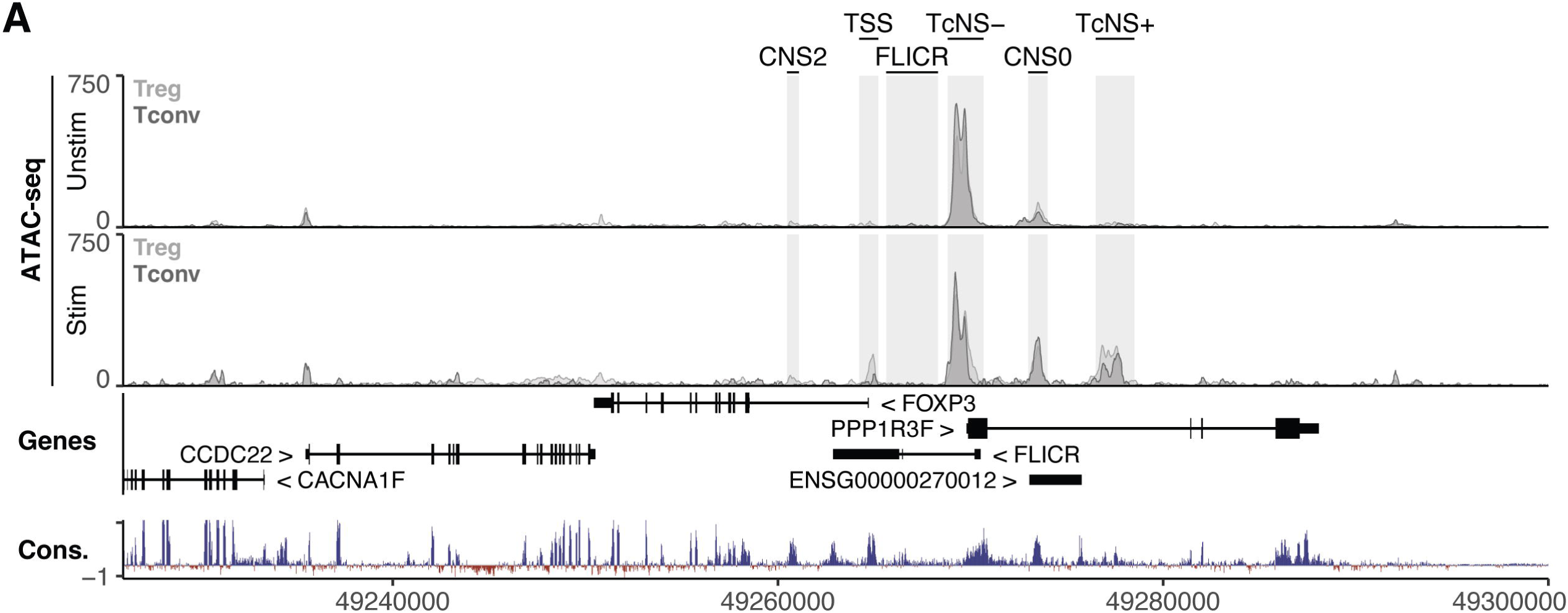
ATAC-seq in stimulated and unstimulated Treg and Tconvs. (A) ATAC-seq in Treg (light gray) and Tconv (dark gray) in unstimulated (top) and stimulated (bottom) conditions^22^.

**Figure S4.**
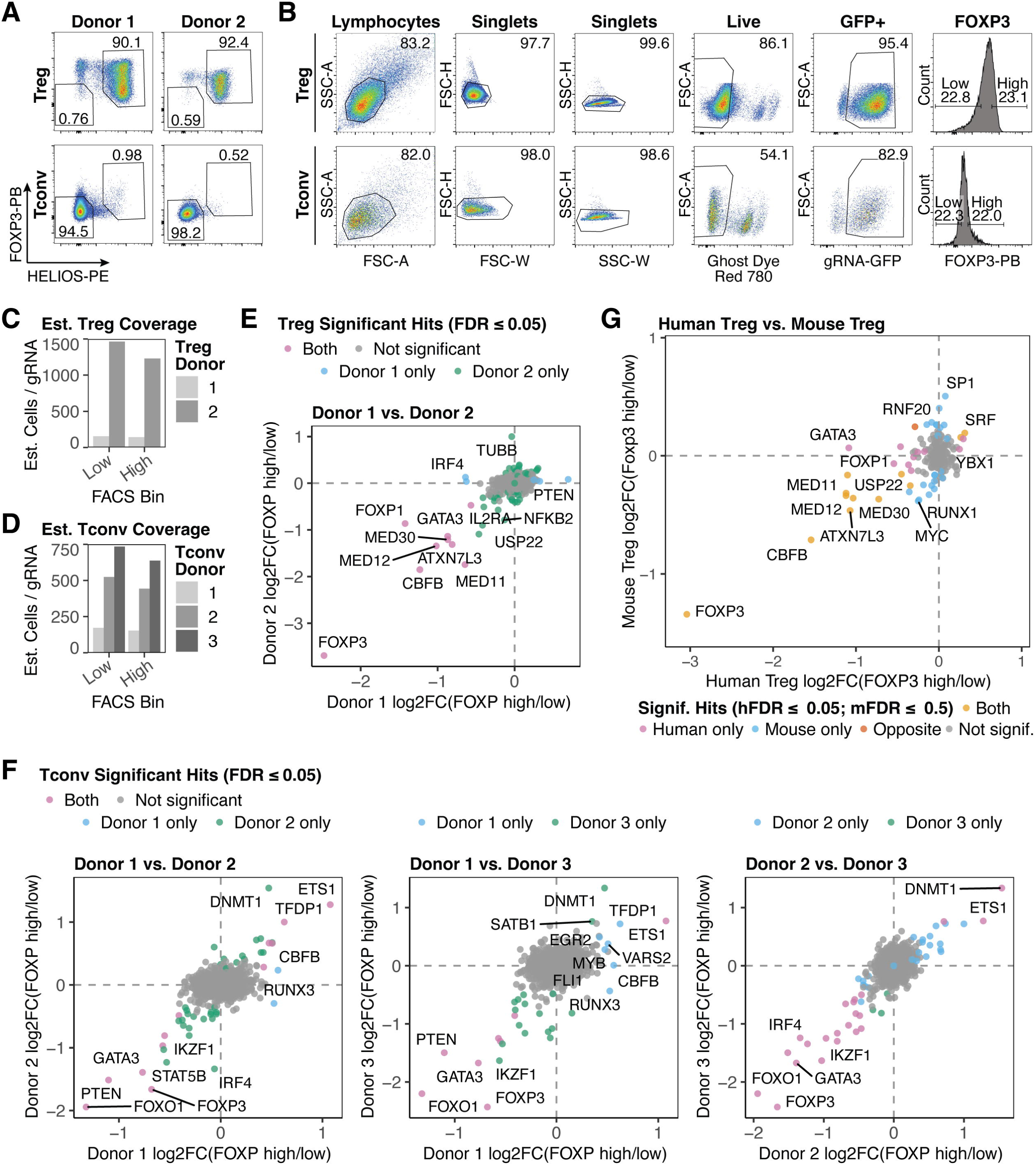
CRISPRn TF KO screens identify FOXP3 trans-regulators. (A) Flow cytometry plots depicting FOXP3 and HELIOS expression in Treg donors used in CRISPRn screens. Donor-matched Tconvs are included for purity comparison. (B) Example representative gating strategy of Treg (top) and Tconv (bottom) FOXP3 expression in CRISPRn tiling screens. (C and D) Estimated cell coverage per gRNA in FOXP3 high and low bins in Treg (C) and Tconv donors (D). Estimated coverage was calculated as the number of sorted cells in each bin divided by 6000, the theoretical number of gRNAs in the trans-regulator library. (E and F) CRISPRn trans-regulator screen donor correlation plots of log2 fold change of gene gRNA enrichment in FOXP3 high vs. low FACS bins in Tregs (E) and Tconvs (F). Color of points indicates significance. Pink, significant in both donors. Blue/green, significant in indicated donor only. Gray, not significant (FDR ≤ 0.05). (G) CRISPRn trans-regulator screen correlation plots comparing log2 fold change of gene gRNA enrichment in FOXP3 high vs. low FACS bins from human Treg screens and a published murine screen^29^. The FDR cutoff for murine significance was FDR ≤ 0.5, as reported in (ref.^29^). Color of points indicates significance. Yellow, significant in both species. Pink, significant in human Tregs only. Blue, significant in murine Tregs only. Orange, discordant effects between human and murine Tregs. Gray, not significant (Human FDR ≤ 0.05, murine FDR ≤ 0.5).

**Figure S5.**
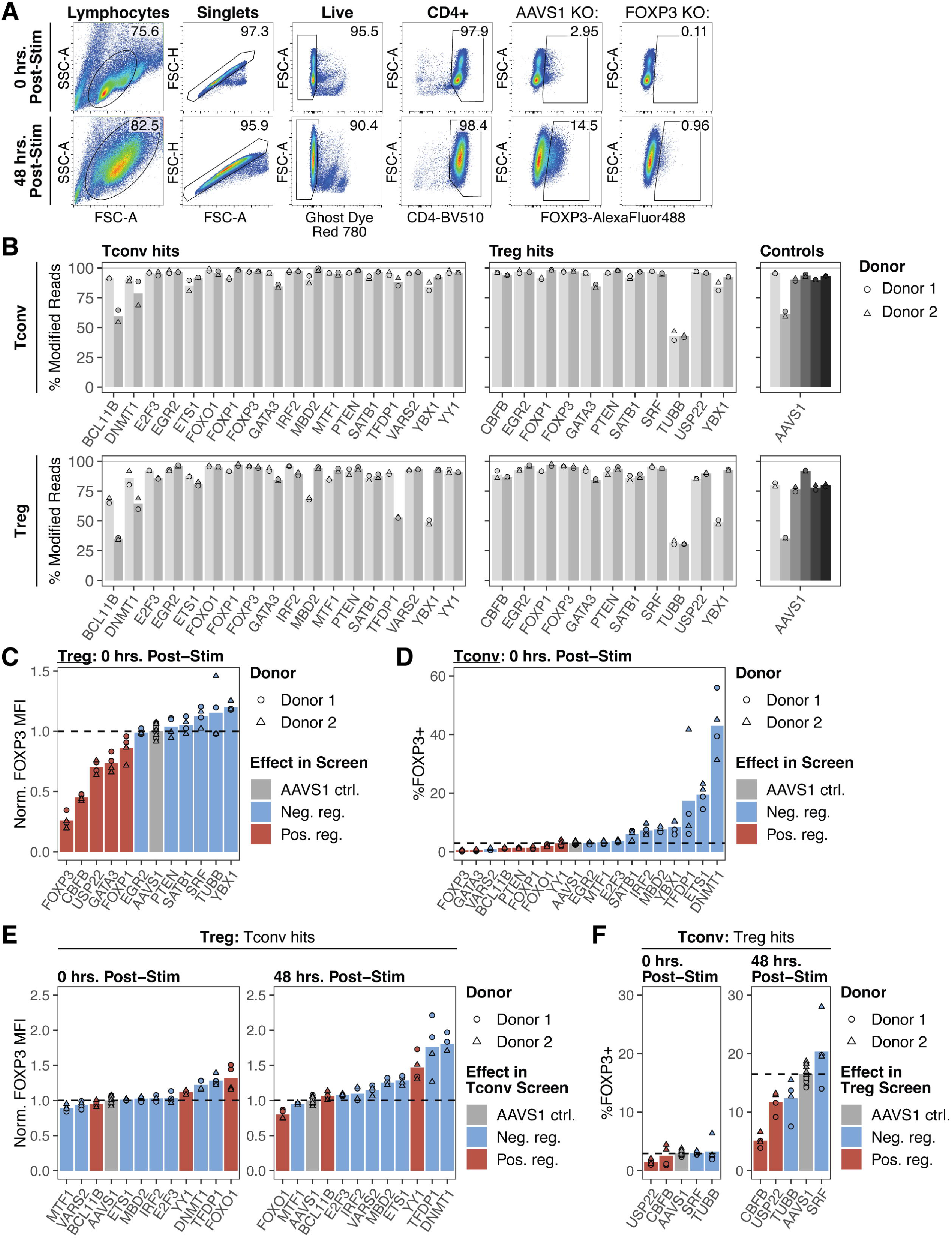
CRISPRn screen arrayed validation characterizes FOXP3 trans-regulators. (A) Representative gating strategy of Tconv %FOXP3+ cells in arrayed validation at 0 hours post-stimulation (top) and 48 hours post-stimulation (bottom). FOXP3+ gate was set on cells receiving a FOXP3 KO gRNA, as shown. (B) Editing efficiency (% modified reads) of arrayed validation genes and *AAVS1*-targeting controls in Tconvs (top) and Tregs (bottom). Color of bars indicates individual gRNAs (*n* = 2 donors x 2 gRNAs per gene or 6 gRNAs targeting *AAVS1* control). (C and D) Arrayed validation of top FOXP3 maintenance and suppressive regulators in Tregs (C) and Tconvs (D) at 0 hours post-stimulation. Color indicates directional effect in screen. In Tregs, FOXP3 MFI in each targeted sample is normalized to the donor-matched mean of *AAVS1*- targeting controls (*n* = 2 donors x 2 gRNAs per target or 6 gRNAs targeting *AAVS1* control). (E) Arrayed validation of unique Tconv FOXP3 maintenance and suppressive regulators in Tregs at 0 and 48 hours post-stimulation. Color indicates directional effect in Tconv screen. FOXP3 MFI for each targeted sample is normalized to the donor-matched mean of *AAVS1*-targeting controls (*n* = 2 donors x 2 gRNAs per target or 6 gRNAs targeting *AAVS1* control). (F) Arrayed validation of unique Treg FOXP3 maintenance and suppressive regulators in Tconvs at 0 and 48 hours post-stimulation. Color indicates directional effect in Treg screen (*n* = 2 donors x 2 gRNAs per target or 6 gRNAs targeting *AAVS1* control).

**Figure S6.**
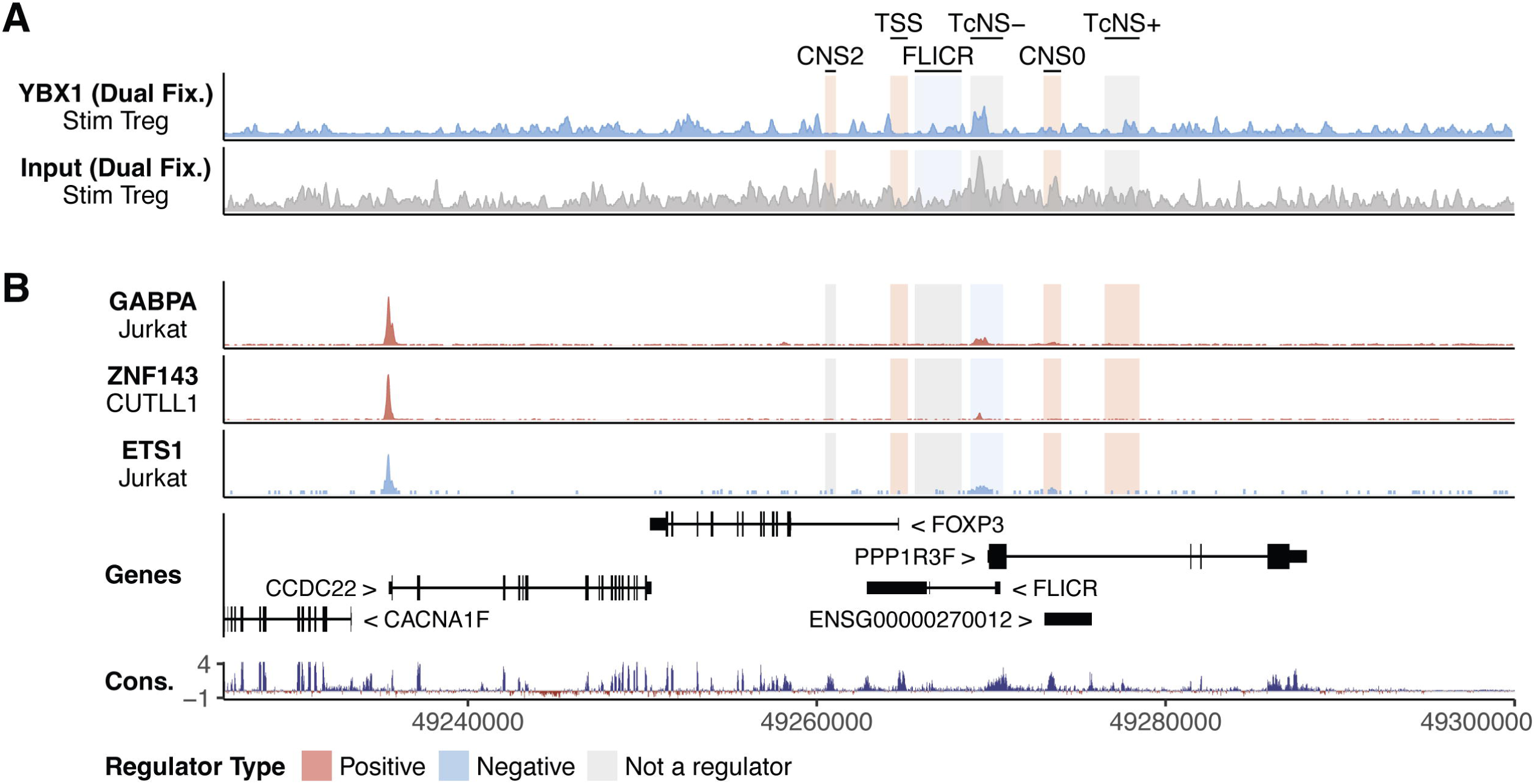
Additional ChIP-seq tracks for FOXP3 trans-regulators. (A) YBX1 (and input) ChIP-seq in stimulated Tregs using formaldehyde and DSG dual crosslinking. ChIP-seq tracks are auto-scaled to the broader FOXP3 locus (chrX:48958000- 49318000). (B) ChIP-seq for GABPA, ZNF143, and ETS1 in T cell lines. ChIP-seq tracks are auto-scaled to the locus shown.

**Figure S7.**
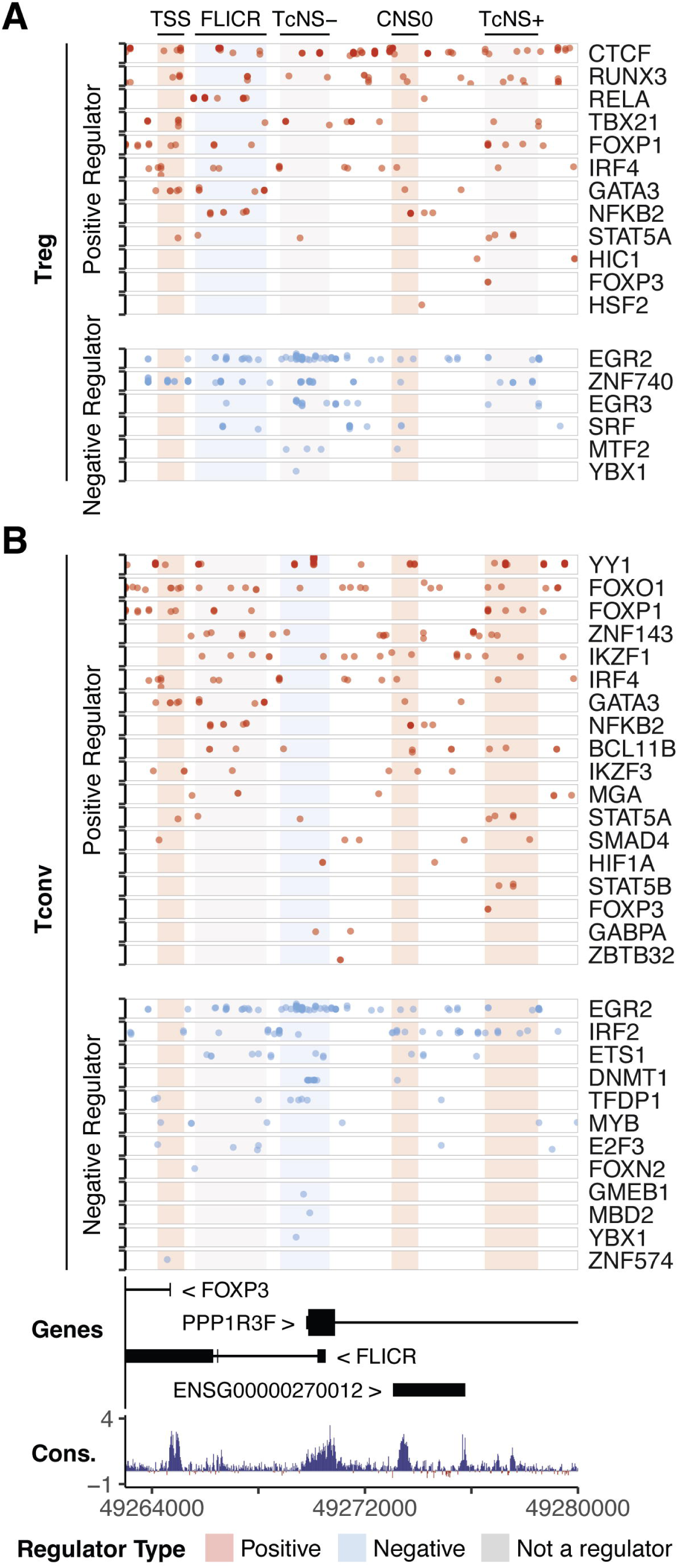
Predicted TF binding motifs of FOXP3 trans-regulators. (A and B) Predicted TF binding sites of positive FOXP3 trans-regulators (top, red) and negative FOXP3 trans-regulators (bottom, blue) plotted along the *FOXP3* locus in Tregs (A) and Tconvs (B). CREs are colored based on the effect of the element in each cell type. Blue, negative regulator; red, positive regulator; gray, not a regulator.

**Figure S8.**
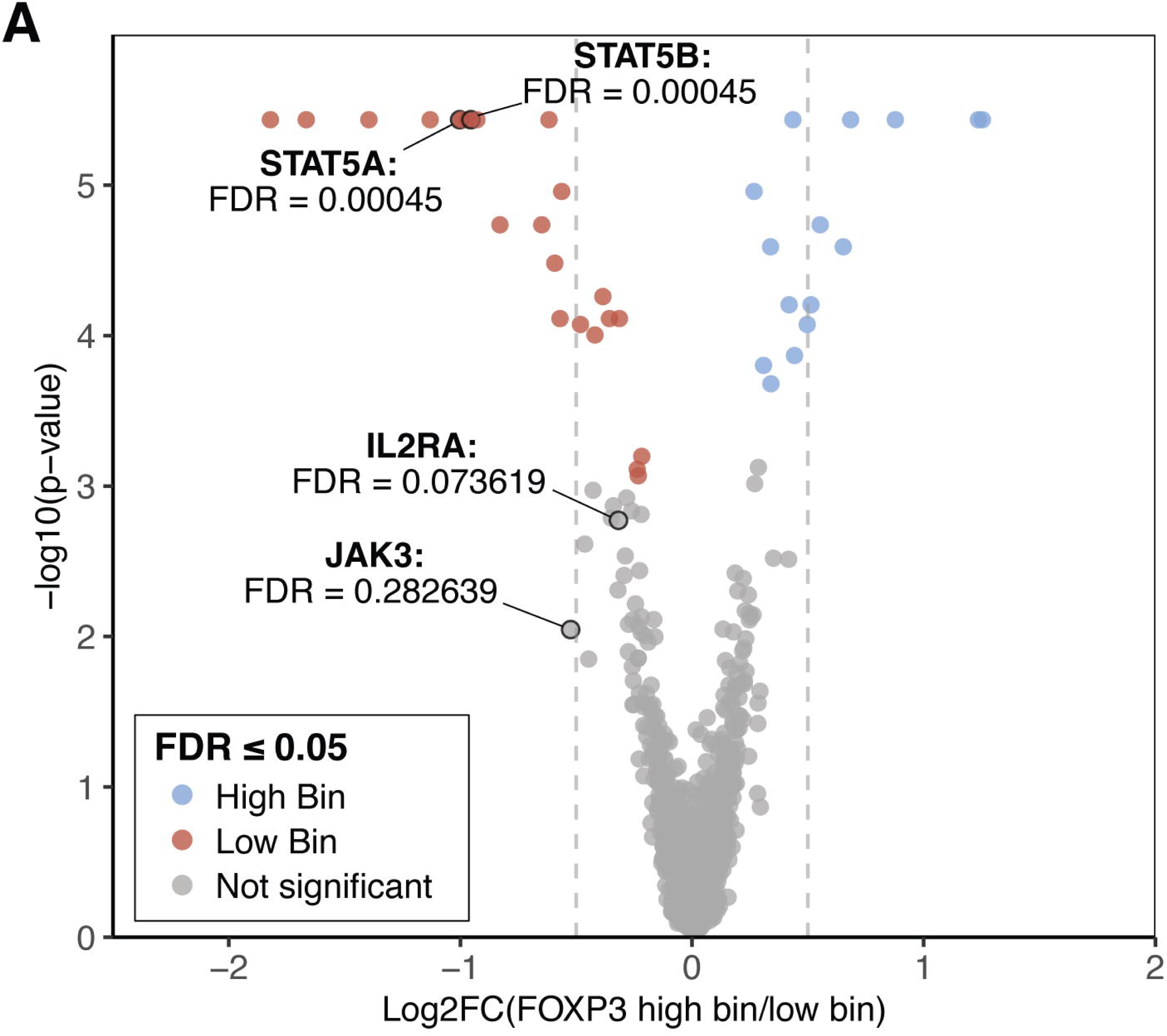
Effect of IL-2 pathway components in Tconv FOXP3 trans-regulator screen. (A) Volcano plot of log2 fold change of gene gRNA enrichment in FOXP3 high vs. low FACS bins versus -log10 of p-value in Tconvs. IL-2 pathway components are labeled with gene name and FDR, and points are indicated with a black border. Color of points indicates significance (FDR ≤ 0.05). Blue, significantly enriched in FOXP3 high FACS bin. Red, significantly enriched in FOXP3 low FACS bin. Gray, not significant (*n* = 3 donors).

**Figure S9.**
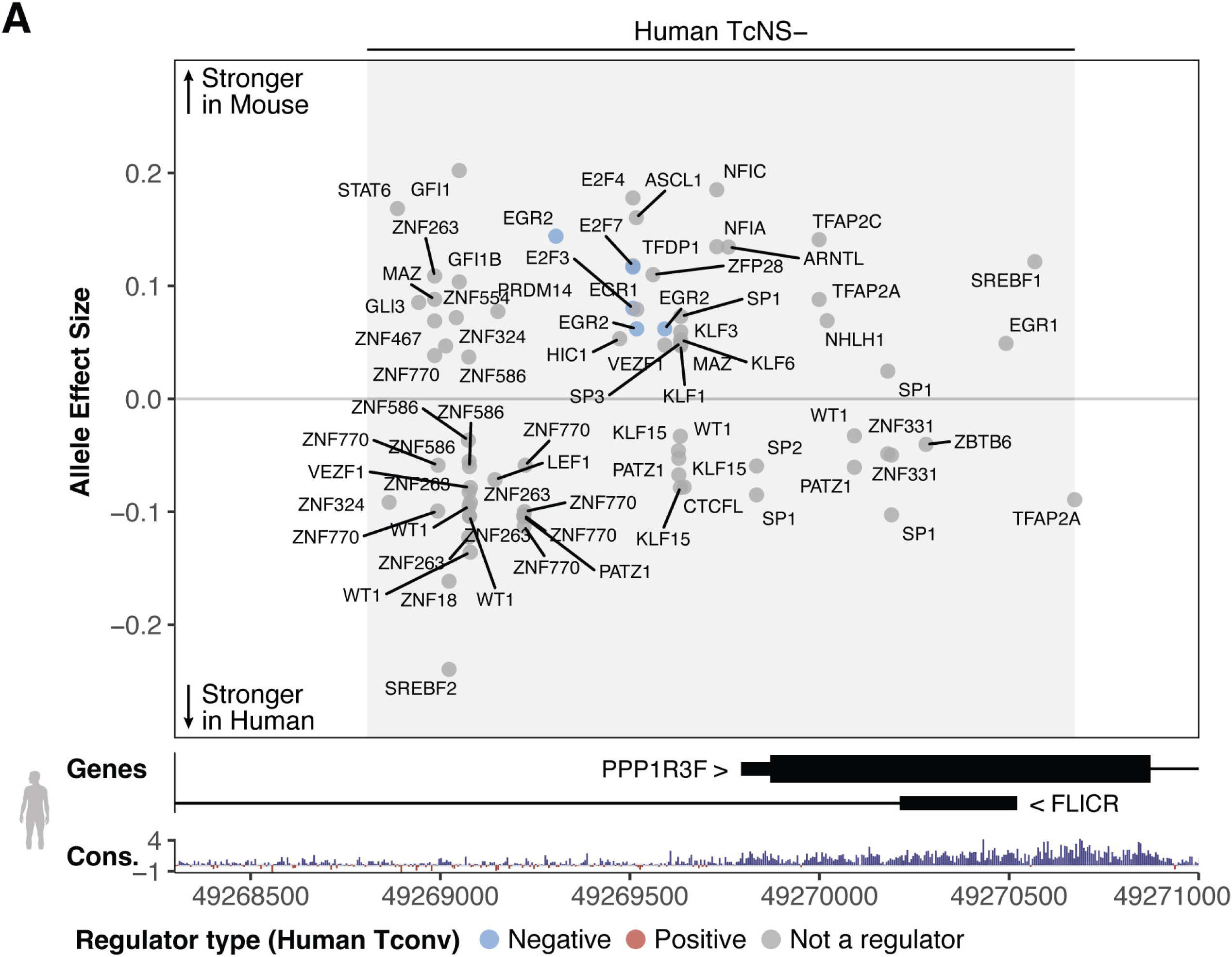
Motif affinity analysis comparison between human and mouse TcNS-. (A) Allele effect size of TF binding motifs between human (reference allele) and mouse (alternative allele) TcNS-, plotted along the human locus. Allele effect size indicates the difference in motif scoring between the reference and alternate allele (alternate – reference)^71^.

**Figure S10.**
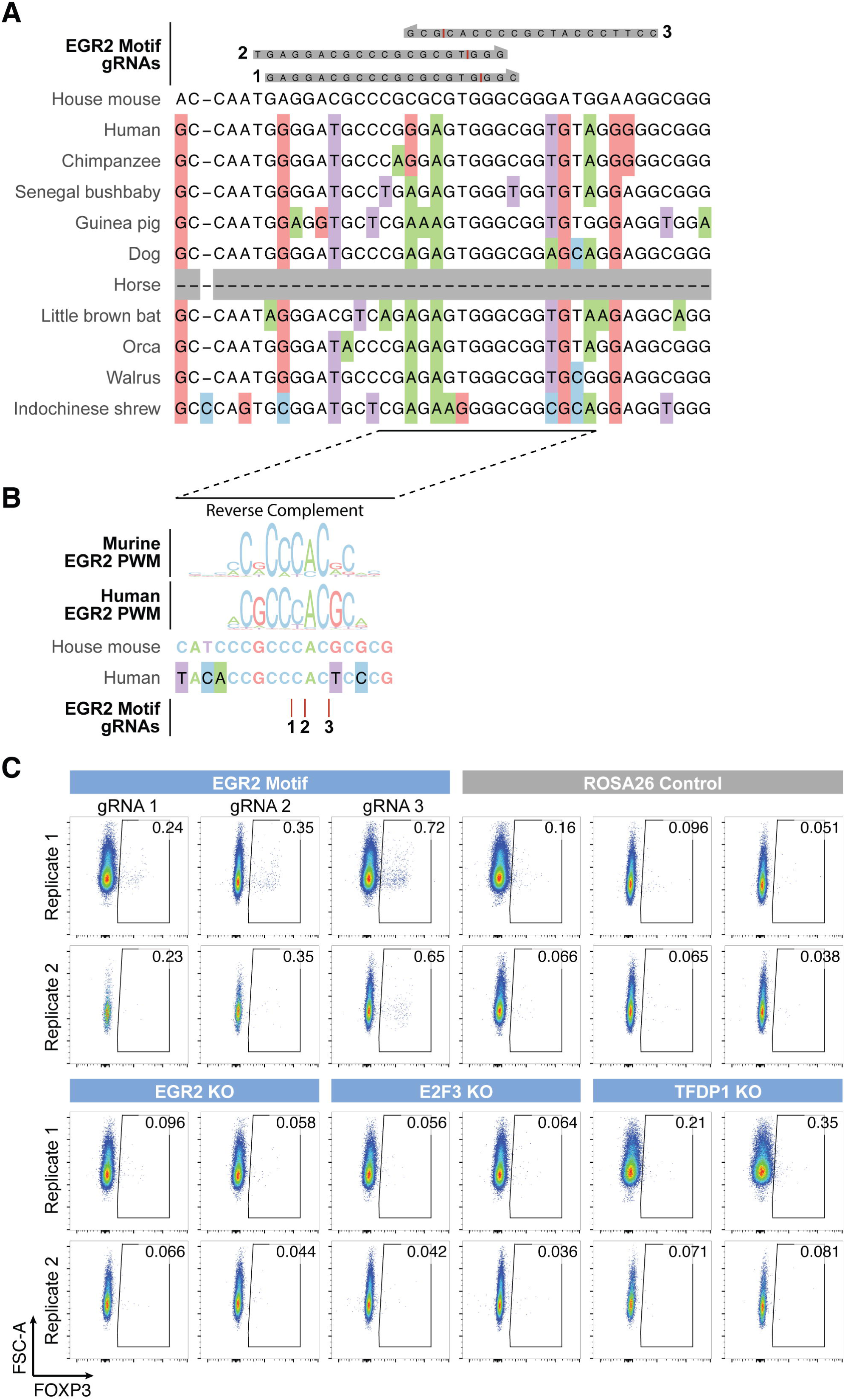
CRISPRn targeting of differential affinity EGR2 motif and associated TFs. (A) Top, gRNAs targeting EGR2 motif. Cut site is indicated in red. Bottom, multiple species alignment of mouse sequence chrX:7574460-7574500 (mm10). Colored boxes indicate deviations from murine reference sequence. Dashes indicate indels between species. (B) Reverse complement of indicated region in panel (A), overlayed with human and murine EGR2 motif sequence logos^75^ (top), sequence comparison (middle), and EGR2 motif cut sites (bottom). Colored boxes indicate deviations from the murine reference sequence. (C) Effect of human negative regulator KO, EGR2 motif disruption, and *Rosa26* Control targeting on FOXP3 expression in murine Tconv 9 days post-stimulation.

